# Evidence for shallow cognitive maps in schizophrenia

**DOI:** 10.1101/2024.02.26.582214

**Authors:** Ata B. Karagoz, Erin K. Moran, Deanna M. Barch, Wouter Kool, Zachariah M. Reagh

## Abstract

Individuals with schizophrenia can have marked deficits in goal-directed decision making. Prominent theories differ in whether schizophrenia (SZ) affects the ability to exert cognitive control, or the motivation to exert control. An alternative explanation is that schizophrenia negatively impacts the formation of cognitive maps, the internal representations of the way the world is structured, necessary for the formation of effective action plans. That is, deficits in decision-making could also arise when goal-directed control and motivation are intact, but used to plan over ill-formed maps. Here, we test the hypothesis that individuals with SZ are impaired in the construction of cognitive maps. We combine a behavioral representational similarity analysis technique with a sequential decision-making task. This enables us to examine how relationships between choice options change when individuals with SZ and healthy age-matched controls build a cognitive map of the task structure. Our results indicate that SZ affects how people represent the structure of the task, focusing more on simpler visual features and less on abstract, higher-order, planning-relevant features. At the same time, we find that SZ were able to display similar performance on this task compared to controls, emphasizing the need for a distinction between cognitive map formation and changes in goal-directed control in understanding cognitive deficits in schizophrenia.

## Introduction

Impairments in goal-directed behavior are a critical aspect of schizophrenia (Barch and Dowd 2010; Cooper et al. 2019; Culbreth et al., 2016; Liemburg et al., 2015). These impairments become prevalent before the onset of psychosis, stabilize afterwards (Green, 1996; Sheffield, Karcher, & Barch, 2018), and are resistant to treatment even when other symptoms are ameliorated (Green, 2016; Tripathi, Kar, & Shukla, 2018). Moreover, they predict long-term and daily life outcomes (Kring & Barch 2014; Cowman et al. 2021), such as a reduced lifespan and a higher likelihood of homelessness (Ayano, Tesfaw, & Shumet, 2019). Understanding how schizophrenia affects goal-directed behavior is a critical step towards developing more effective diagnosis and treatment. However, the mechanistic underpinnings of these impairments remain poorly understood.

Most prior research has explained these deficits in goal-directed behavior as the result of a reduced capacity for cognitive control (Barch, Culbreth, & Sheffield, 2017). For example, Cohen and Servan-Schreiber (1993) argued that disturbances of dopaminergic function reduce the quality of information processing in the prefrontal cortex, an area of the brain critical for cognitive control (Miller & Cohen, 2001). This may lead to a global reduction in the ability of patients to perform the computations needed to plan towards goals (Culbreth et al., 2016; Knolle et al., 2023). This might occur through a reduced ability to manipulate information in working memory (Kim et al. 2004), to maintain relevant information (Thakkar and Park 2010; Gotra, Keedy, and Hill 2022) or to suppress distraction (Reilly et al. 2008; Lesh et al. 2011). Another line of research argues that deficits in goal-direct control instead arise from perturbed estimations of mental effort demands (Gold et al., 2013; Gold et al., 2015), potentially driven by abnormal dopaminergic functioning (Westbrook et al., 2021). Under this explanation, reduced goal-directed control in schizophrenia reflects a decision to withhold mental effort (Cooper et al., 2019; Culbreth, Westbrook, & Barch, 2016) rather than an inability to apply it. In short, both theories focus on how a decrease in implemented control produces cognitive deficits in schizophrenia, only differing in whether capacity or perceived incentive is the source of disruption.

However, deficits in goal-directed behavior can also arise if one exerts appropriate control over ill-formed representations of the world (Feher da Silva & Hare, 2020; Feher da Silva et al., 2023). Representations of the world are key to effective goal-directed decision-making as they provide maps as to the stimuli and/or actions that could lead to varying outcomes. If individuals with schizophrenia are not able to generate accurate representations of the work, goal-directed decision-making impairments would manifest even if individuals with SZ were motivated and able to expend effort on a given task. That is, as long as internal models, or cognitive maps (Tolman, 1948), inaccurately reflect the structure of the world, any good faith attempt to plan or reason over them will result in performance deficits. Thus, deficits in goal-directed control in schizophrenia may occur because patients have difficulty constructing models of the environment.

There is some preliminary evidence to suggest that schizophrenia interferes with the ability to construct appropriate internal representations of task structure. For example, individuals with schizophrenia often fail to integrate intrinsically linked characteristics of choice options (Morris et al. 2018; Cooper et al., 2019), need to be explicitly pointed to task structure to perceive differences in effort demands between choice options (Gold et al., 2015), and show blunted internal error monitoring signals (Kirschner & Klein, 2022). Individuals with schizophrenia are also impaired at combining previously learned associations to form new inferences (Armstrong, Williams, and Heckers, 2012), navigating complex virtual environments (Weniger and Irle, 2008), and updating internal representations of dynamically changing task rules (Everett et al. 2001; Waltz and Gold 2007; Morris et al. 2018). Many of these deficits center on relational processing (Einstein & Hunt, 1980; Halford et al., 2010), which underlies a host of cognitive abilities ranging from decision-making (Abbasov et al., 2011) to episodic memory functions mediated by the hippocampus (Davachi, 2006; Shimamura, 2011; Whittington et al., 2022). Importantly, the hippocampus is thought to be critical for the formation of cognitive maps of the world around us (O’Keefe and Nadel 1978; Behrens et al. 2018). This set of findings about cognitive impairments have led to the recently proposed “shallow cognitive map” hypothesis of schizophrenia. This hypothesis claims that these relational deficits stem from disorganization in hippocampal circuity, leading to disorganized thought (Musa et al. 2022). If mechanisms such as those in the hippocampus underlying the formation of cognitive maps (O’Keefe & Nadel, 1978; Whittington et al., 2022) are disrupted in schizophrenia, then many deficits in schizophrenia – including goal-directed decision making – may be a consequence of poorly-built internal representations of the world. However, this hypothesis has not been directly tested, especially in contexts in which people need to use goal-directed control to plan over their internal representations of a task’s structure.

We test this hypothesis by leveraging a recent formalization of goal-directed control in terms of “model-based” reinforcement learning (RL; Sutton & Barto, 2018). Model-based decision-makers search through their internal representation of the environment to find the lines of action that produce the maximal cumulative reward. This form of decision-making is often contrasted with “model-free” reinforcement learning, which simply updates the value of actions that lead to reward without considering the structure of the task (Daw et al., 2005; Kool, Cushman, and Gershman, 2017; Drummond & Niv, 2020). It has been argued that human behavior reflects a weighted mixture of these two strategies. This RL framework of goal-directed behavior provides a computationally explicit distinction between control and the representations over which control is applied. The latter is a function that links actions to consequences (a cognitive map), whereas the former is an algorithm that uses this function to search for rewards.

We have recently developed a novel behavioral approach that measures people’s subjective representation of a task structure that they need to navigate to earn rewards (Karagoz, Reagh, & Kool, 2024). At the heart of this approach lies a variant of a class of sequential decision-making tasks, called two-step tasks, that dissociate model-based from model-free control (Daw et al., 2011; Kool et al., 2016). This paradigm is paired with a behavioral representational similarity analysis (behRSA) approach (Karagoz, Reagh, & Kool, 2024). We developed this analysis to use simple similarity judgments between objects encountered in the task to assess how participants represent the structure of the task (inspired by neural variants of this same analysis; Kriegeskorte et al., 2008). Importantly, this approach allows us to decompose participants’ cognitive maps into three increasingly complex components that represent distinct forms of structure in the two-step task. These components range from representing structure that is irrelevant to task performance to the most complex higher-order structure that is beneficial for planning. In our prior work, (Karagoz, Reagh, & Kool, 2024), we found that participants whose cognitive maps better aligned with higher-order structure in the task tended to use more model-based control, and performed better overall. Based on the shallow cognitive map hypothesis, we predicted that schizophrenia patients would construct mental representations of the task that overweight simple surface-level features and underweigh the more complex higher-order relationships that are relevant for planning.

In order to test for motivational deficits in goal-directed control implementation, we included a “stakes” manipulation in this task, so that on certain trials the earned reward would be multiplied. This manipulation temporarily amplifies the benefit of model-based control, and results in a phasic increase in its use in conventional research populations (Kool et al., 2017; Karagoz, Reagh, & Kool, 2024; Patzelt et al., 2018). This shift has been argued to result from a cost-benefit tradeoff that pits the computational costs of planning against its increased accuracy (Kool & Botvinick, 2018; Kool et al., 2017). Even though schizophrenia appears not to affect reward processing (Heerey, Bell-Warren, & Gold, 2008; Abler et al., 2008), it should still affect sensitivity to amplification of rewards if effort costs are higher (Kring & Barch, 2014; Treadway et al., 2015; Reddy et al., 2015; Cooper et al., 2019; Barch et al., 2023). For instance, patients might still integrate reward values but consistently deem the cost of effort to not be worth the increase in reward. The stakes manipulation allowed us to test this hypothesis.

Our results provide novel evidence for the shallow cognitive map hypothesis of schizophrenia. The cognitive maps of patients tended to focus on features of the task that are not relevant to planning but are perceptually salient, whereas controls reported higher similarity for some higher-order planning-relevant features. Interestingly, in this simple version of the two-step task, we found that patients and controls performed approximately equally well to one another. Taken together, this provides important evidence that model-based control and cognitive map construction are separable constructs, and that internal models of task structure are indeed disrupted in patients with schizophrenia. Finally, we found that patients were not sensitive to motivational manipulations, providing additional evidence that schizophrenia affects the motivation to exert goal-directed control.

## Methods

### Participants

We recruited 20 people with schizophrenia/schizoaffective disorder (SZ), and 23 control participants (CN) to participate in the study. Exclusion criteria included: 1) DSM-5 diagnosis of substance abuse or dependence in the past 6 months; 2) IQ less than 70 as measured by the Wechsler Test of Adult Reading (Wechsler 2001); and 3) history of severe head trauma and/or loss of consciousness. Additional exclusion criteria for patient group included inpatient or partial hospital status. Additional criteria for controls included: 1) no personal or immediate relative with a history of schizophrenia, schizoaffective disorder and 2) no current or past major depression. Finally, we also excluded participants based on failing to respond to more than 20% of trials in the decision-making task (a single control participant). All participants provided written informed consent to the protocol approved by the Washington University Institutional Review Board. Demographics for all groups are presented in Table 1. There were no group differences in either age, sex or parental education, though as is typical, the individuals with schizophrenia had slightly lower personal education.

**Table 1:** Participant demographic characteristics.

|  | Healthy Controls (N=23) |  | Individuals with Schizophrenia (N=20) |  |  |
| --- | --- | --- | --- | --- | --- |
|  | Mean | SD | Mean | SD | p-value |
| **Demographics** |  |  |  |  |  |
| Age (years) | 38.85 | 8.91 | 42.82 | 9.98 | 0.18 |
| Sex (% male) | 52.17% |  | 55.0% |  |  |
| Personal Education (years) | 15.30 | 2.70 | 13.35 | 2.81 | 0.03 |
| Parental Education (years) | 13.15 | 2.80 | 14.14 | 3.03 | 0.29 |
| **Self-Report** |  |  |  |  |  |
| Snaith-Hamilton Pleasure Scale | 12.26 | 1.96 | 10.45 | 3.00 | 0.03 |
| Motivation and Pleasure Scale | 38.48 | 10.67 | 31.65 | 13.06 | 0.07 |
| **Neurocognitive Measures** |  |  |  |  |  |
| Wechsler Test of Adult Reading | 103.43 | 8.84 | 102.10 | 13.57 | 0.71 |

### Diagnostic and symptom assessment

Diagnostic status was confirmed using the Structured Clinical Interview for DSM-5 conducted by masters or Ph.D.-level clinicians. Clinician-rated negative symptoms were assessed in all patient groups using the Clinical Assessment Interview for Negative Symptoms (CAINS)(Kring et al. 2013), which includes a Motivation and Pleasure (MAP) and Expression (EXP) subscale. General psychiatric symptoms were assessed using the Brief Psychiatric Rating Scale (BPRS)(Overall and Gorham 1962) which includes a subscale of psychotic symptoms and depression. Participants completed the Motivation and Pleasure Scale (MAP-SR; Llerena et al. 2013) with higher scores equaling more motivation and pleasure across the week. Participants also completed the Snaith-Hamilton Pleasure Scale to assess hedonic capacity with higher scores equaling more capacity (Snaith et al. 1995). Depression was assessed using the Center for Epidemiological Studies Depression Scale (CES-D; Radloff 1977).

### Decision-making task

The decision-making task was designed to dissociate between model-free and model-based decisions in a setting where participants need to learn about the structure of the task and was based on a recently developed ‘two-stage’ sequential decision-making task (Kool, Gershman, and Cushman 2017; Karagoz, Reagh, and Kool 2024).

Each trial of the task started randomly in one of two first-stage states. Each of these states offered a choice between a unique pair of ‘teleporters’, presented side-by-side. Participants used the ‘F’ key on their keyboard to choose the left teleporter, and the ‘J’ key to choose the right teleporter. This choice determined which one of two second-stage states would be encountered. For each pair, one of the teleporters deterministically led to a purple second-stage state, and the other deterministically led to a red second-stage state. Importantly, each teleporter always led to the same second-stage state (Figure 1A).

**Figure 1.**
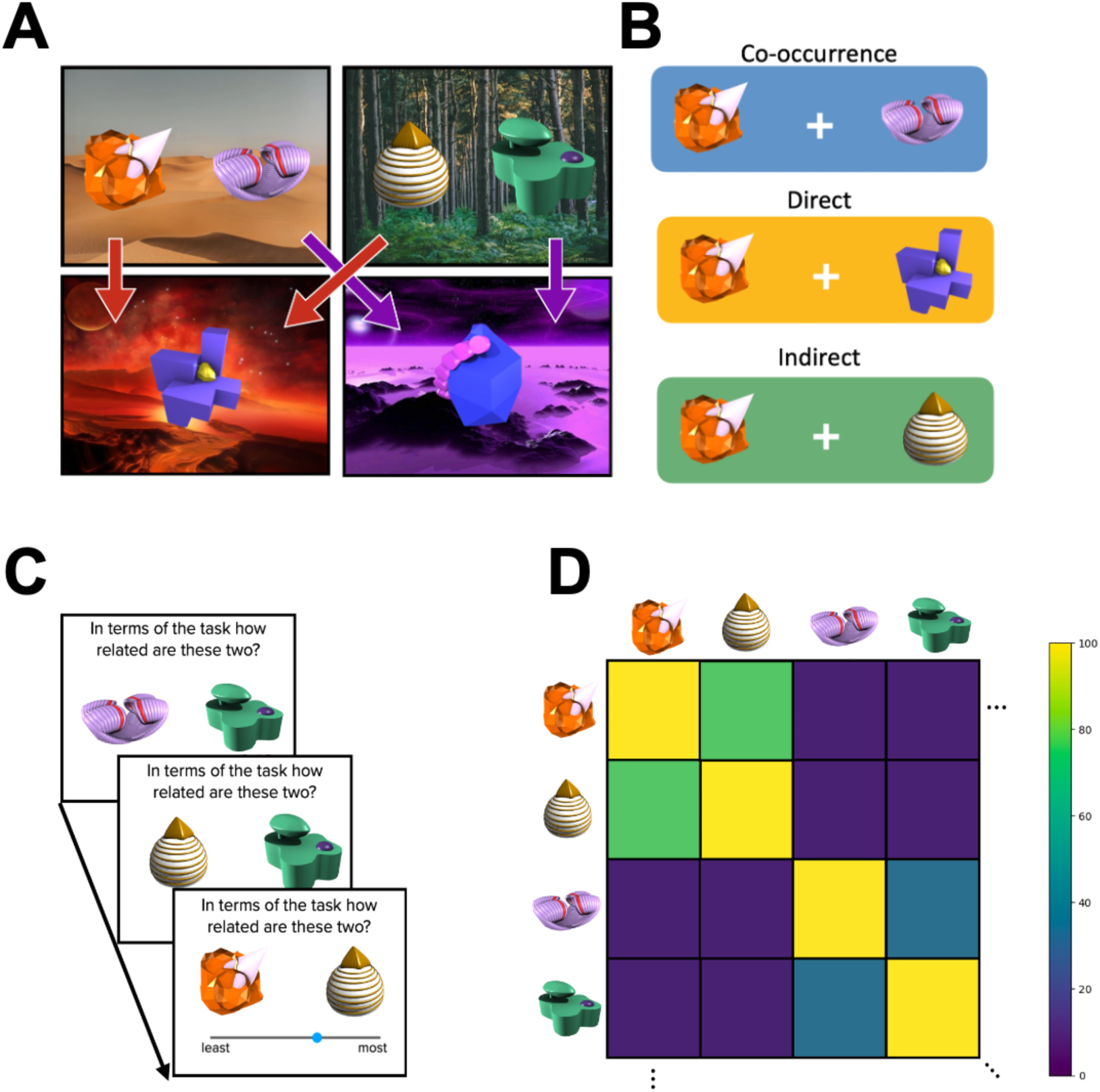
Task schematic. **A.** Task transition structure. two distinct first-stage each contain two unique choice objects. Each of these objects deterministically leads to one of two second-stage states, as depicted by the colored arrows. These first-stage states were associated with different amounts of reward that changed across the task. **B.** Possible relationships between items that can be inferred from task experience. First, items that appear together can become more subjectively related (co-occurrence). Second, items that form an action outcome pair can become more subjectively related (direct). Third, items that share a goal state can become more subjectively related (indirect). **C.** Example trials from the relatedness task. Participants see a pair of objects they experienced in the context of the decision-making task and are asked to move a slider to indicate the degree of ‘relatedness’. **D.** A portion of a hypothetical matrix of similarity ratings generated from the behRSA task. behRSA = behavioral representational similarity analysis

Each second-stage state contained a unique “generator” that was associated with a scalar reward. Participants were instructed to press the spacebar key to interact with the generator so that it provided them with “space treasure”, and they were told that the fuel rods used by the generators would sometimes yield more or less space treasure. The payoffs of the generators changed over the course of the experiment according to independent random walks. Their reward distributions were initialized randomly for each participant within a range of 0 to 9 points and then varied according to a Gaussian random walk (*σ* = 2) with reflecting bounds at 0 and 9.

The 6 objects were randomly assigned as teleporters and generators for each participant separately.

This task distinguishes between model-based and model-free strategies, since only a model-based decision maker generalizes experiences from one starting state to all other starting states. That is, after receiving a high reward in a second-stage state, a model-based learner can use their knowledge of the transition structure to plan their way back to that same second-stage state. A model-free agent, on the other hand, learns through action-reward associations and will only become more likely to choose that same action in the same first-stage state, not transferring experiences from one first-stage state to the others (Kool, Cushman, and Gershman 2016). If we imagine a trial starting in the desert first-stage state that leads to a better-than-expected reward in the red second-stage state, a purely model-free agent will not use this information if the next trial starts in the forest first-stage state. The computational model, described below, uses reinforcement-learning to capture this distinction in a single model-based weighting parameter.

In order to introduce differing incentives for model-based control, we introduced a ‘stakes’ manipulation in this task (Kool, Gershman, and Cushman 2017). At the start of each trial, an incentive stake cue indicated by how much the reward obtained at the end of the trial would be multiplied. On some trials, this cue indicated that the points would be multiplied by 5 (high stakes). On other trials, the cue indicated that the points would be multiplied by 1 (low stakes). For example, if a participant earned 5 space treasure pieces on a high-stakes trial, the multiplier would result in a total of 25 points. On a low-stakes trial with the same amount of space treasure, the participant would earn 5 points. The chance of a high stakes trial was equivalent for both first-stage states. On each trial, there was a 50% chance that a trial would be a high stakes trial and a 50% chance it would be a low stakes trial.

At the start of each trial, participants saw the first-stage background and the stake multiplier for 1 second. Then, the stake moved to the top left corner and the teleporters were presented. Participants were then given a time limit of 3 seconds to choose between them. After their response, the selected option was highlighted, and the non-selected option was greyed out for the remainder of the response period. There was a 500ms interval between the end of the first stage response period and the onset of the second stage. Following a 200ms interval after the generator was selected, the space treasure pieces produced by the generator were displayed at the top of the screen for 1.5 seconds. Each piece was individually converted to a point value (100ms for each) and then the points for that trial were multiplied by the stake before being added to the score in the top-right. There was a 500ms intertrial interval (ITI). Participants completed a total of 200 trials with an optional short break in the middle.

### Behavioral representational similarity task

To test how task experience affects structure learning, we used a behavioral representational analysis technique previously described by Karagoz and colleagues (2023). In this task, participants provided relatedness ratings of pairs of novel 3D objects that were also used as choice options in the decision-making task (adapted with permission from (Hsu, Schlichting, and Thompson-Schill 2014; Schlichting and Preston 2015). On each trial, they were shown two of the objects experienced in the task horizontally centered in the screen (image sizes of 400 x 400 pixels). At the top of the screen the prompt read “In terms of the task, how related are these two objects?”. Below the images, participants were shown a ‘slider’ bar and were asked to use the mouse to move the slider to their perceived level of ‘relatedness’ between the two objects they had just seen (Figure 1C). Participants had unlimited time to respond, with a 1 second inter-stimulus interval between submission and the next relatedness trial. Participants performed pairwise ratings of all 6 objects in both left and right positions, resulting in 30 trials.

We have used this approach prior work (Karagoz, Reagh & Kool 2024), where we have shown that participants’ relatedness ratings reveal their assessment of task structure. For example, a participant may think that a first-stage teleporter and the second-stage reward generator it leads to, are related, while the same teleporter is not related to the generator on the other second-stage state. As we detail below, our analytic method allows us to quantify these correspondences using a-priori components.

### Working memory task

Participants completed a Running Span task to assess working memory. Letters were presented on a computer screen one at a time, spaced 2-s apart. During a trial, an unpredictable number of letters was presented, and participants were asked to remember the last x number of letters from the list. The task began with participants recalling the last letter presented and progressed in difficulty in each successive block of 4 trials. Participants had to complete 2 out of 4 trials correctly to move on to the next block. The dependent measure was the total number of correct letters recalled in correct order across all trials.

### Procedure

Participants completed one visit to the laboratory as part of a larger study examining motivation and cognition. Participants completed a diagnostic interview followed by tasks assessing reinforcement learning, working memory and questionnaires assessing symptom domains as described above. Participants were compensated $40 for their time and for any task bonus they received.

### Analysis

#### Performance

We computed average performance on the decision-making task as the average number of points earned per trial. To correct for baseline differences in available reward (as a result of the random Gaussian walks), we then subtracted the average available reward across both second-stage states. We also computed the average performance based on the stake effects. For this, we took the difference between participant performance in high-stakes vs low-stakes trials to indicate a measure of motivational modulation of performance.

#### Reinforcement learning model

We adapted an established hybrid reinforcement learning model that we used in prior work to assess participants’ behavior in the decision-making task, specifically dissociating model-free and model-based decision making (Kool et al. 2016; Bolenz et al. 2019; Karagoz, Reagh, and Kool 2024). Full details of the model and fitting procedure are discussed in the Supplementary Methods.

After each action, the model-free system computes a prediction error, which is simply the difference between the current value (both future and immediate reward) and the reward expectation. It then uses a temporal difference-learning algorithm (Sutton & Barto 2018) to increase values for actions in response to positive prediction errors and lower values that lead to negative prediction errors. The model-based learner combines the transition structure of the task with second-stage model-free values to plan its first-stage choices, and does not rely on first-stage model-free values.

These two learning systems are combined using a weighting parameter (*w*) bounded between 0 and 1, where 0 is fully model-free control and 1 is fully model-based. The combined system then made choices using a stochastic choice rule (soft-max), using an inverse-temperature parameter β that governs the explore/exploit tradeoff. The model also included a learning-rate ɑ that determines the degree to which prediction errors update existing action values, an eligibility trace parameter λ that controls how the outcome at the second stage informs the first-stage, and finally π and ρ which capture perseveration on either response or stimulus choice.

We used maximum a posteriori estimation to fit this dual-system RL model to behavior on this task, using empirical priors previously reported by Bolenz et al. (2019). The average fit per group for each of these parameters is reported in Table 2.

**Table 2:** Reinforcement learning model parameter estimates. None of the parameters differ significantly between the two groups although there is a trending effect of better model fits in the patient group.

|  | Healthy Controls (N=23) |  | Individuals with Schizophrenia (N=20) |  |  |
| --- | --- | --- | --- | --- | --- |
|  | Mean | SD | Mean | SD | p-value |
| LL | -116.30 | 21.84 | -104.80 | 16.55 | 0.06 |
| $\alpha$ | 0.69 | 0.22 | 0.60 | 0.27 | 0.26 |
| $\beta$ | 1.58 | 1.10 | 1.62 | 1.12 | 0.87 |
| $\lambda$ | 0.58 | 0.09 | 0.60 | 0.18 | 0.86 |
| $\rho$ | -0.09 | 0.29 | -0.06 | 0.40 | 0.80 |
| $\pi$ | 0.25 | 0.60 | 0.56 | 0.78 | 0.16 |
| $w$ | 0.52 | 0.16 | 0.48 | 0.16 | 0.13 |

#### Behavioral Representational Similarity Analysis

We used participants relatedness ratings of objects to measure the structure of their cognitive maps (Figure 1CD). Based on our previous work with this variant of the task, we hypothesized that experience with the decision-making task would yield participants to represent the task structure across three different levels of abstraction (Karagoz, Reagh, and Kool 2024). These levels of abstraction map onto the possible relationships highlighted in Figure 1B. We call the first level of abstraction ‘visual co-occurrence’, which corresponds to relatedness between items observed in the same first-stage state. Even though this relationship between items is valid, it does not aid goal-directed control. The second “direct association” is defined as the relationship between a first-stage transporter and the second-stage generator, and the third highest-order “indirect association” is the relationship between two first-stage teleporters that share the same goal. Though these differ in terms of level of abstraction, both these components are relevant for planning. Using the direct associations allows participants to infer the consequence of a first-stage action, whereas the indirect associations allows them to infer that two items that both lead to the same second-stage state are functionally equivalent. To test this hypothesis, we compared their matrices of relatedness ratings to three a-priori model matrices, which reflected these levels of representation of the decision-making task structure. This was done using a multiple regression approach where the flattened vector of similarity ratings provided by participants was the dependent variable. Each of the a-priori model matrices was also flattened and modeled as an independent variable. The final regression formula was as follows:

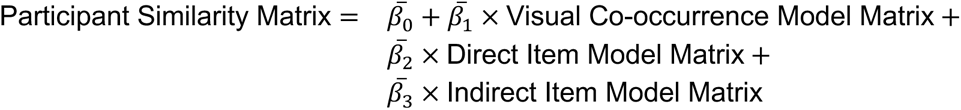

This formulation allowed us to analyze the participant similarity matrix as a weighted average of the different planning relevant and irrelevant features in the task. We used these beta values as further regressors to see if amount of reported similarity along one of our a-priori features predicted either performance or use of model-based control.

#### Hierarchical linear mixed effects model (behRSA)

We modeled the relationship between the inferred task structure, clinical group, and the degree of complexity of task structure features using a hierarchical linear mixed effects model. Specifically, we model edthe estimated regression coefficients described in the previous section as a function of categorical regressor that encoded clinical group (patient vs. control) and an ordinal regressor that encoded the complexity of the component (co-occurrence = −1, direct = 0, indirect = 1), while including random intercepts for each participant.

#### Linear model for effect of self-report data on *w*

We modeled the relationship between use of model-based control, clinical group, and the self-report scores from both the Snaith-Hamilton Pleasure Scale and the Motivation and Pleasure Scale (MAP-SR) using a linear model. Before incorporating the raw scores into the model, we first z-scored each self-report measure within group. These values were then used in the linear model with a binary indicator of clinical group where control is 0 and patient is 1.

#### Statistical approach

In our statistical analyses, we relied on two-tailed tests wherever prior research did not inform a predicted direction of the effect. We used one-tailed tests when previous results from our group allowed us to make directional predictions about effects. When running independent samples t-tests between our groups we use Welch’s tests.

## Results

### Differences in task behavior

Inconsistent with prior research on schizophrenia and model-based control, we found that the patients performed the task in a qualitatively similar way to controls. Patients and controls did not differ in terms of chance-corrected reward rate *t*(30.91) = −0.28, *d* = 0.09, *p* = 0.779) (Figure 2A). One intuitive hypothesis for this finding is that patients spent more time on first-stage choices, but we found no group difference in first-stage response times (*t*(38.16) = −1.04, *d* = 0.32, *p* = 0.304). We also assessed to which degree patients and controls failed to respond on a trial. Numerically, the control group showed a smaller average percentage of missed trials (mean % missed = 0.04 in controls and 0.06 in patients), but this difference between group non-responses was not significant (*t*(30.23) = −1.30, *d* = 0.41, *p* = 0.203).

**Figure 2.**
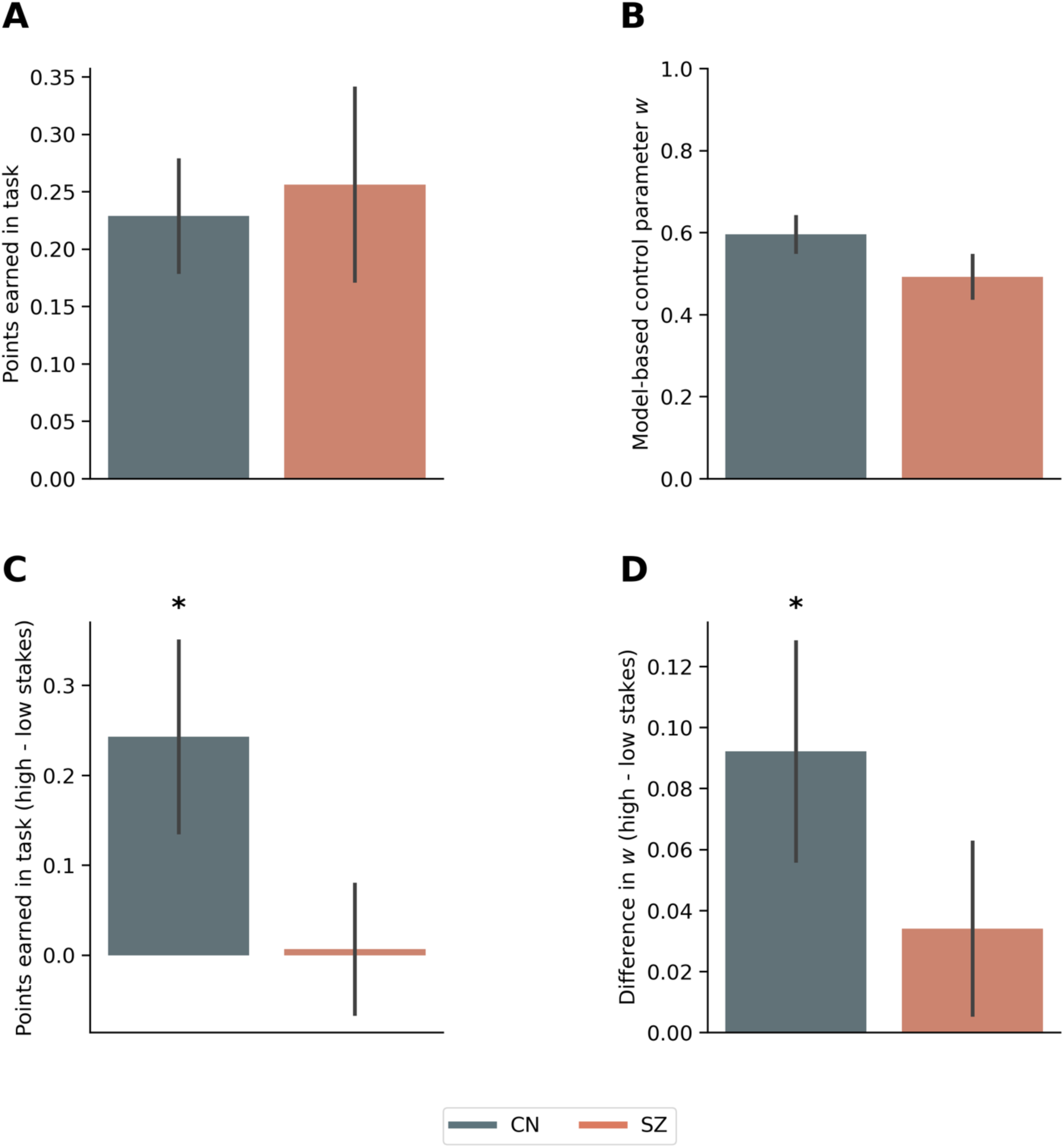
Differences in task performance. **A.** Baseline adjusted points earned during the task. **B.** Model-based control parameter *w* **C.** Differences in points earned by stake condition. **D.** Differences in model-based control by stake condition. (* indicates p < 0.05, error bars are standard error of the mean)

The computational model fits provided consistent results. In contrast to previous findings, we saw no difference in the model-based control parameter between the two groups (*t*(38.55) = −1.55, *d* = 0.48, *p* = 0.129) (Figure 2B). One interesting possibility is that this is driven by the fact that increased use of model-based control in our task leads to an increased payoff (Kool et al., 2016), whereas the original version of this (Daw et al., 2011) used in prior work (Culbreth et al., 2016) does not. Another explanation is that model-based control in this task is less demanding because it does not require reasoning over probabilistic transitions (unlike the original version). In short, these results suggest that patients were able and willing to engage in control over goal-directed behavior.

Next, we turned our attention to the incentive manipulation. Prior work has shown that people increase model-based control when higher reward is available (Kool et al. 2017; Bolenz et al. 2019; Karagoz et al., 2023). In our study, we found that we find in healthy controls an increase in points earned in the high-stake trials compared to the low-stake trials (*t*(22) = 2.29, *d* = 0.48, *p* = 0.032), but patients showed no such sensitivity to the stake manipulation (*t*(19) = 0.09, *d* = 0.02, *p* = 0.926) (Figure 2C). More specifically, patients had similar means in both stake conditions (µ_high_ = 0.263, µ_low_ = 0.256), while controls differed in mean points stakes (*μ*_high_ = 0.344, *μ*_low_ = 0.101) (Figure S2). The computational modeling fits were consistent with this pattern of behavior. For controls, we saw an effect of stake on model-based control with higher stakes resulting in higher model-based control (*t*(22) = 2.57, *d* = 0.54, *p* = 0.017), but we did not see this effect in patients (*t*(19) = 1.21, *d* = 0.27, *p* = 0.242) (Figure 2D).

### Differences in cognitive maps

In order to test the shallow cognitive map hypothesis of schizophrenia, we analyzed the data from the behRSA task. In previous work, this task allowed us to characterize participants’ cognitive maps across three independent components, increasing in complexity. We found that the more complex components predicted task performance (Figure 1B, direct and indirect item associations) (Karagoz, Reagh, and Kool 2024). Here, we used this approach to assess whether patients and controls differ in the way they construct cognitive maps.

To do so, we ran a hierarchical linear mixed-effects with participants’ model matrix coefficients as the dependent variable and both clinical group (patients vs. controls) and the complexity of the component (co-occurrence = −1, direct = 0, indirect = 1) as regressors. There was no significant main effect of clinical group, (*β* = 6.40, *CI*95% = [−3.1,15.9], *p* = 0.18) or component complexity (*β* = −1.9, *CI*95% = [−13.3,9.5], *p* = 0.74). However, there was a significant interaction between clinical group and component complexity (*β* = −14.6, *CI*95% = [−28.6, −0.7], *p* = 0.039), indicating that cognitive maps of schizophrenia patients underweighted the more abstract and complex features of the task structure (see Figure 3).

**Figure 3.**
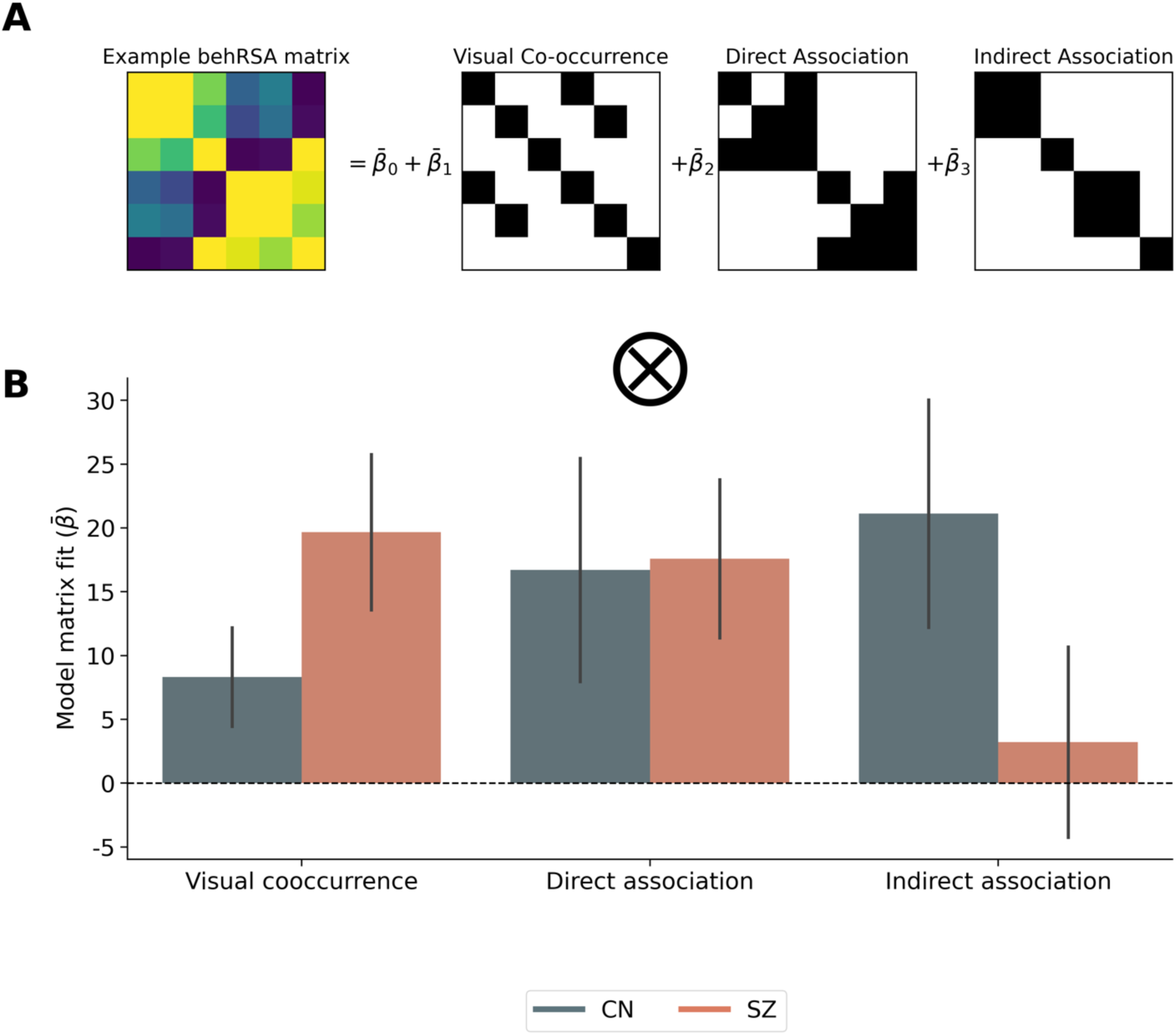
Differences in cognitive map building. **A.** Behavioral representational similarity analysis approach. Participants’ post-task judgments of relatedness are turned into a pairwise similarity matrix. The lower triangle of this matrix is then compared to three model matrices corresponding to 3 hypotheses about features participants may use in their mental representation. **B.** Patients show a heightened sensitivity to visual co-occurrence, rating it quantitatively higher than control participants. The two groups show approximately equal weighting of direct associations that link first-stage items to their second-stage counterparts. Patients seem to not use the indirect association between the two first-stage items that lead to the same second-stage item. Controls report this feature in keeping with previous work. (Error bars are standard error of the mean; ⦻ indicates a significant interaction between group and fit across model matrices.)

Next, we aimed to assess whether patient and control representations contain group consistent structure that is not accounted for by our model. Therefore, we calculated the Spearman rank correlation between the lower triangle values of the behRSA matrix of a given participant and the mean values for all other participants within their group. We then repeated this, leaving each participant out once, until we had an estimate for the intra-class correlation of the behRSA representations. Interestingly, patients had a relatively high mean intra-class correlation (average Spearman rho = 0.252, std = 0.255). Control participants meanwhile had a lower average intra-class correlation value, with more variance (average Spearman rho = 0.129, std = 0.377). Thus, individuals with schizophrenia were more similar to each other when performing the relatedness task. Control participants, on the other hand, were more varied. Given that several pairwise relationships between items could be identified (as is seen in our prior work; Karagoz, Reagh, & Kool, 2024), this could indicate that patients were more constrained in the set of relationships that they represented in their cognitive maps.

After investigating these group differences of the principal measures of our tasks, we turned our attention towards individual differences. We first focus our analyses on individual differences in model-based control, and then report a set of analyses that explore individual difference in cognitive map formation.

### Individual differences in model-based control

#### Basic effects

The two-step decision-making task that we used in this study was designed in such a way that increased model-based control produces increased rewards. Indeed, previous work using variants of this task have confirmed this through robust correlations between the degree of model-based control, as measured by the *w* parameter, and points earned in the task (Kool et al., 2016; Bolenz et al. 2019; Karagoz et al., 2023).

In our control sample, we replicate this relationship (*r*(21) = 0.55, *CI* 95% [0.18,0.79], *p* = 0.006) (Figure 3A in blue). Interestingly, we do not find a significant correlation between patient use of model-based control and reward gained in the task (*r*(18) = 0.08, *CI* 95% = [−0.38,0.5], *p* = 0.741) (Figure 3A in orange). One potential explanation for this difference is that patient choice was more exploratory or random, but we found no difference in the inverse-temperature parameter (*β*) between groups, (see Table 2). We also examined whether differences in the explore/exploit tradeoff (as indicated by *β*) accounted for the difference in correlations between model-based control and performance. However, we found no evidence that controlling for individual variation in the inverse-temperature parameter uncovered a relationship between *w* and points earned for patients (Supplementary Table 1).

Importantly, other key RL parameters from our model fits predicted performance in both groups. For example, differences in the inverse temperature positively predicted points earned in both the control group, (*r*(21) = 0.70, *CI* 95% = [0.40,0.86], *p* < 0.001), and the patients, (*r*(18) = 0.55, *CI* 95% = [0.14,0.8], *p* = 0.012). Analogously, individual differences in the learning rate parameter predicted performance in the control group,(*r*(21) = 0.63, *CI* 95% = [0.29,0.83], *p* = 0.001), and in patients (*r*(18) = 0.63, *CI* 95% = [0.26,0.84], *p* = 0.003). These results confirmed the general predictive validity of our RL modeling approach.

#### The effect of stakes incentives

Next, we examined the correlations between the benefits of heightened control and increased performance in the high-stakes context. In the control group, we found that the degree to which control is heightened in high stakes trials positively predicts the degree of increased performance (*r*(21) = 0.59, *CI* 95% = [0.23,0.8], *p* = 0.003). Meanwhile, in the patient group we found no such benefit of modulation of control (*r*(18) = 0.30, *CI* 95% = [−0.17,0.66], *p* = 0.200) (Figure 4B).

**Figure 4.**
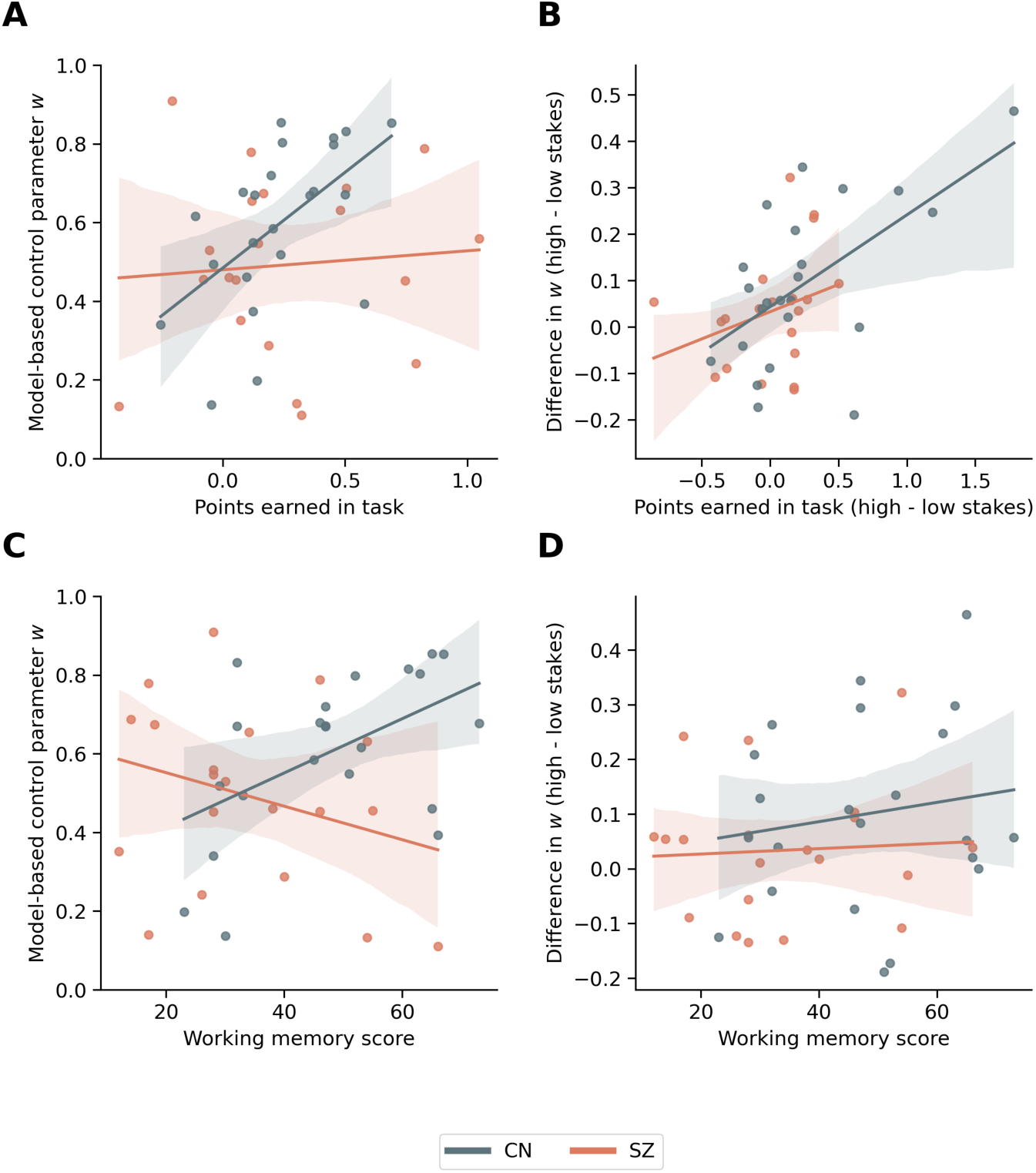
Correlations between model-based control and performance. **A.** Controls, but not patients, show a strong correlation between model-based control and task performance. **B.** The amount to which control participants increase their model-based control in high-stakes contexts is correlated with increased performance in those contexts. This relationship is absent in patients. **C.** Working memory correlates strongly with model-based control in healthy controls, but not in patients. **D.** Working memory does not predict increases in model-based control in response to the stakes manipulation.

#### Working memory

Next, we investigated task correlations with the working memory scores that we measured with the Running Span task. It is important to note that due to missing data from a control participant the working memory results have 22 controls and 20 individuals with schizophrenia. In line with previous work (Culbreth et al. 2016; Otto et al., 2013), we found a significant relationship between use of model-based control and working memory in healthy controls (*r*(21) = 0.51, *CI* 95% = [0.11,0.77], *p* = 0.015), but we found no such correlation for patients’ (*r*(18) = −0.29, *CI* 95% = [−0.65,0.18], *p* = 0.219). We also assessed whether working memory capacity was coupled with the increase of model-based control on high-stakes trials, but we found no relationship between modulation of control and working memory in controls (*r*(21) = 0.16, *CI* 95% = [−0.28,0.54], *p* = 0.489) or patients (*r*(18) = 0.06, *CI* 95% = [−0.39,0.49], *p* = 0.801). As noted under Basic Effects above, we found no relationship between model-based control and reward earned in patients. We examined whether this lack of relationship in patients relative to controls could be explained by differences in working memory capacity. Modeling Running Span as a covariate in a group-wise regression analysis, we found that working memory differences did not explain decoupling of model-based control from reward in patients (see Supplemental Materials, Table S7).

Finally, we assessed correlations between use of model-based control and our measures of motivation and pleasure seeking as well as hedonic capacity. We found that, within both groups, hedonic capacity as measured by the Snaith-Hamilton Pleasure Scale and motivation as measured by MAP-SR were positively correlated (*r*(41) = 0.33, *CI* 95% = [0.03, 0.57], *p* = 0.033). To assess how these self-reported measures interact with use of model-based control we ran a linear model predicting the effect of each along with interactions with clinical group. Our measure of hedonic capacity did not predict the use of model-based control in healthy controls (*β* = 0.057, *CI* 95% = [−0.063,0.176], *p* = 0.342). In previous work, Culbreth and colleagues (2016) found a positive relationship between an estimate of model-free control and hedonic response. Here, we replicate this finding. In our model, where lower *w* values indicate more model-free control, we found that use of model-free control was again correlated with higher hedonic capacity (but only in patients). More specifically, we found a negative relationship between hedonic capacity and model-based control in patients (*β* = −1.62, *CI* 95% = [−0.314, −0.010], *p* = 0.037). Motivation and pleasure-seeking scores did not predict model-based control in healthy controls (*β* = −0.079, *CI*95% = [−0.181,0.022], *p* = 0.123). Interestingly, we observed a trending interaction where motivation and pleasure-seeking predict a relative increase in use of model-based control in patients (*β* = 0.126, *CI*95% = [−0.017,0.269], *p* = 0.082). We further ran models assessing how self-report measures predicted other features of our task such as modulation of control, these are reported in the Supplemental Results: MAP-SR and Snaith-Hamilton section.

### Individual differences in cognitive map formation

In our previous work, we found that participants who had stronger representations of the direct item and indirect item associations showed increased task performance (Karagoz, Reagh, and Kool 2024).

Even though our current sample size was considerably smaller than in our previous study, we found modest evidence that these effects replicate. Due to our previous results, we ran one-tailed tests here for positive correlations. In controls, we found a correlation between task performance and the strength of the direct item associations (*r*(21) = 0.40, *CI* 95% [0.05, 1.0], *p* = 0.030), but we found no such effect for indirect item associations (*r*(21) = 0.18, *CI* 95% [−0.18, 1.0], *p* = 0.206, both of these were one-tailed tests based on our prior work). In patients, representation of the direct item association showed no relationship with task performance (*r*(18) = 0.18, *CI* 95% [−0.28, 0.58], *p* = 0.437), neither did the indirect item association (*r*(18) = 0.14, *CI* 95% [−0.32, 0.55], *p* = 0.544). We next assessed whether components of the behRSA were associated with differences in the use of model-based control. In the present data, we found no indication in either our controls or patients that model-based control was coupled with behRSA components.

#### Correlation of cognitive map components and stake effects

Due to the fact that we saw differences in control modulation of *w* between high and low stakes conditions we ran a model to examine whether modulation of control was predicted by the presence of different associations in participant model matrices. That model did not account for any variance in the data (Supplement: *w* modulation by behRSA).

We then ran a model to look if differences in points earned were predicted by the subject behavioral representations. We found that there was a difference in the degree to which direct item associations predicted the modulation of reward by stakes. In the healthy controls, there was a positive relationship between the presence of direct item associations and the degree to which more points were earned in higher-stakes (*β* = 0.009, *CI*95% = [0.003,0.015], *p* = 0.005). Meanwhile, we observed an interaction indicating that this relationship is not present in patients (*β* = −0.015, *CI*95% = [−0.015, −0.004], *p* = 0.01). Interestingly, we found a trending opposite effect in the more abstract indirect item association. In control participants, increased reporting of indirect associations was linked to less modulation of performance (*β* = −0.006, *CI*95% = [−0.012,0.000], *p* = 0.056). Interestingly, in patients, the presence of indirect item associations in the cognitive map was linked to increase performance modulation (*β* = 0.010, *CI*95% = [0.001,0.019], *p* = 0.035). These results indicate major differences in the way control participants and patients represented components of the task, and how these representations were used to guide behavior. In particular, patients’ representations of the task do not assist them in modulating their effort to increase reward. (full results of the model are displayed in Table 3).

**Table 3:**
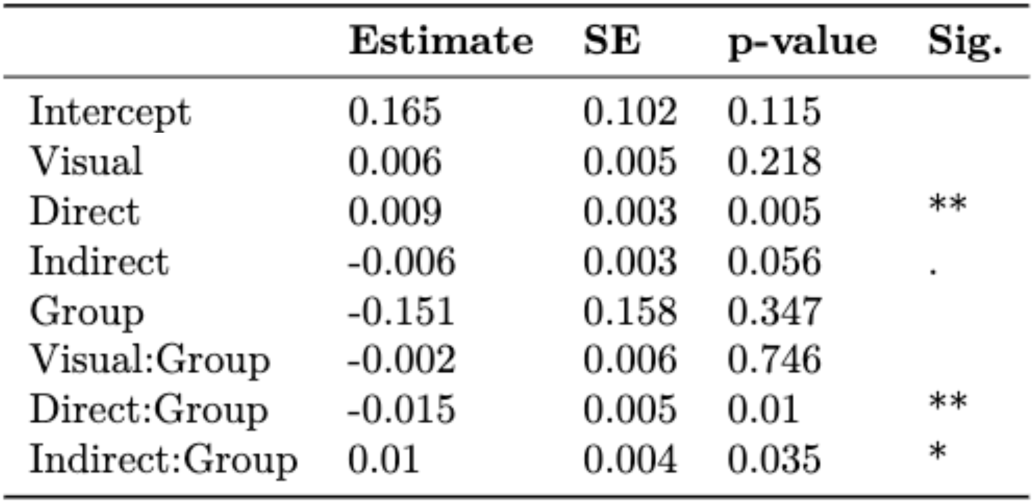
Differences in performance between the stake manipulation as predicted by behRSA fits.

## General Discussion

Patients with schizophrenia often show marked deficits in goal-direct behavior (Barch and Dowd 2010). These deficits are typically understood as an inability or a reduced willingness to exert control (Barch, Culbreth, & Sheffield, 2017; Culbreth, Westbrook, & Barch, 2016; Knolle et al. 2023). Here, we report evidence that suggests that such deficits may also arise because of a reduced ability to construct representations of the environment. Making use of a novel behavioral representational similarity approach, we found that patients construct cognitive maps of a decision-making task that favor simpler, easily inferable, relationships and that undervalue higher-order ones that are more relevant to planning. In addition, the cognitive maps of patients were more uniform with one another (patients had higher intra-class correlation) than those of controls, suggesting patients are more uniform in representing the task. Interestingly, these simpler and less optimal cognitive maps in patients did not lead to a reduction in goal-directed control. While this may seem surprising at first, we found that the cognitive maps of patients reliably represented simple but planning-relevant relationships between actions and subsequent states, which are enough to optimally perform the task. This dissociation provides unique evidence for the idea that schizophrenia affects map formation independently of impairments in the implementation of control. In line with previous work, we also found that patients are unable or unwilling to modulate their degree of effort in exchange for increased payoffs (Gold et al. 2013, Cooper et al. 2019). Together, these results indicate that deficits in goal-directed control in schizophrenia are multifaceted, emphasizing the need for precise behavioral and computational tools to distinguish between multiple contributions.

Our framework is an example of this approach. Specifically, it allows for simultaneous measurement of model-based control and the representations that such control is exerted over (Karagoz, Reagh, and Kool 2024). As noted above, we found relatively intact model-based control, but a deterioration of more complex representations of task structure. These differences in the representations of task structure were coupled with a general increase in uniformity for the cognitive maps reported by patients. These findings are consistent with and provide novel evidence for the recent shallow cognitive map hypothesis of schizophrenia (Musa et al. 2022). This theory posits that dysfunction of the hippocampus in patients results in a shallowing of ‘attractors’, stable states of activity that represent distinct components of memory representations. This shallowing, or instability, of attractors would then cause spurious associations between various features of cognitive maps. In our case, these spurious associations could underlie two key observations in the patients’ behavior. First, their tendency to perceive *all* items in the task as being broadly more similar to one another than controls. Second, their tendency to regard items that simply occurred together in the task as being highly related, even though such items in fact led to opposite outcomes. Relatedly, one could hypothesize that this shallowing of cognitive maps is more likely to affect higher-order relationships that rely on relationships across multiple pairs of stimuli. In future work. our novel approach can be combined with neuroimaging to more directly measure the change in hippocampal representations posited by the shallow cognitive maps hypothesis, and with computational modeling using neural networks to formalize how schizophrenia affects the encoding of structural information (Sučević and Schapiro 2023).

Previous work from our group has reported reductions in model-based control in individuals with schizophrenia compared to healthy controls (Culbreth et al. 2016). While we find that model-based control is quantitatively lower in patients, this difference was not significant in the present data. Several important factors may explain this difference. First, in contrast with the task used by Culbreth et al. (2016; Daw et al., 2011), the two-step task reported here is designed to reward model-based control (Kool et al. 2016). Therefore, individuals with schizophrenia may have been more motivated to use model-based control. Second, goal-directed control may simply be less demanding in our task, as it does not require participants to reason over stochastic transitions. Indeed, young children show hints of model-based control on a similar variant of this simpler task, but not on the more complicated original version (Smid et al., 2023). In sum, individuals with schizophrenia might have less difficulty exerting control here than in the task used by Culbreth and colleagues (2016). Thus, the current two-step task provides an excellent tool for studies of goal-directed control in schizophrenia. Moreover, our findings introduce the possibility that reductions in model-based control found by Culbreth and colleagues (2016) are driven by ill-formed cognitive maps. This hypothesis could be tested using our behRSA approach (Karagoz, Reagh, and Kool 2024).

This interpretation of our data is complicated by the surprising finding that patients do not show a correlation between the degree to which they exert model-based control and the rewards they earn in the task. This is particularly striking since we have reported such correlations in prior work (Kool et al. 2017, Patzelt et al. 2018, Bolenz et al. 2019, Karagoz, Reagh, and Kool 2024), and we also find this correlation in the control group. Several intuitive explanations for this dissociation come to mind. Perhaps participants behaved more randomly, or learned less about the task, and this reduces the impact of the model-based control parameter on decision making. However, we find no discernable differences between groups in performance on the task. Potentially, the patients in our study are relying on a choice strategy that is not well captured by our RL model. For example, it has recently been suggested that people may rely on “successor representations” when solving RL tasks (Momennejad et al. 2017). These are cached representations of expected future transitions from a given state (i.e., which states typically follow the current one), and stand in contrast with cognitive maps that contain the full transition structure. This form of RL is computationally distinct from the representations used by model-free and model-based choice strategies. Our finding that patients encoded the direct association (linking previous and future states), but not the indirect one (linking first-stage states), provides preliminary evidence for this hypothesis. Future investigations, that allow for the measurement of a less dichotomous set of RL strategies (Collins & Cockburn 2020) may provide a better insight into this puzzle.

The tradeoff between model-based and model-free control has been suggested to be governed by a cost-benefit analysis (Kool and Botvinick 2018; Kool Gershman Cushman 2018). At the core of this claim lies a set of studies that demonstrate that people exert more model-based control when more rewards at stake (Kool et al. 2017, Bolenz et al. 2019, Karagoz, Reagh, and Kool 2024). We replicate these findings in healthy controls, but this motivational modulation of model-based control does not seem to occur in patients. This could be due to patient deficits in rapidly integrating the presence of the stake cue and using it to shift strategies (Cooper et al. 2019). Another possibility is that the modulation of control is not seen as worthwhile as patients previously have been shown to be less sensitive to effort demands (Gold et al. 2015). Regardless of the underlying mechanism, these results are consistent with the idea that patients are less easily motivated to exert cognitive control (Gold et al. 2015; Culbreth, Westbrook, and Barch 2016).

In our previous work, we found a strong coupling of planning relevant features in the cognitive map and task performance (Karagoz, Reagh, and Kool 2024). Our current data partly replicates this finding in a much smaller sample, but also reveals a decoupling in patients. As before, we find that the planning-relevant direct item association correlates with general performance in healthy controls. However, we also found that task representations the modulation in planning in response to the stakes manipulation. In control participants, the direct item associations predicted increased performance in the high-stakes condition. Indeed, if planning is costly, it may be particularly useful to rely on the task structure when rewards are temporarily amplified. Interestingly, this coupling is not found in patients even though they report a similar amount of direct item association as controls. Note that this result mirrors the lack of a correlation between model-based control and performance discussed above. This provides yet further evidence for altered representational frameworks in patients, and once again emphasizes the need for the development of tasks that measure a range of decision-making strategies.

We found that working memory capacity predicted reliance on model-based control in our control sample (Culbreth et al. 2016). This finding is in line with a set of studies that relate cognitive control, RL, and working memory (Otto et al. 2013, Culbreth et al. 2016, Gillan et al. 2016, Collins et al. 2014), which suggest that model-based control critically draws on executive functioning capacities implemented by the prefrontal cortex. In the clinical group, however, this correlation was absent. One possible explanation for this difference, apart from the ones mentioned above, is that working memory performance was generally lower for patients. That is, individual differences may become less reliable when people perform poorly on working memory tasks. Future, higher-powered, replications of this effect will be needed to assess these relationships more definitively.

There are a variety of neural circuits that have previously been implicated in deficits in goal-directed behavior in schizophrenia. First, there is evidence of general dysfunction in the dorsolateral prefrontal cortex (DLPFC) (Barch and Ceaser 2012), which may reduce patients ability to exert control in pursuit of goals. However, previous work that has found DLPFC dysfunction in schizophrenia has not separately examined the task representations that patients use. Therefore, it could be that this apparent dysfunction of DLPFC is instead driven by this region attempting to exert control over ill-formed maps, leading to patients abandoning these attempts after failure. Second, the hippocampal formation is thought to be crucial for the construction of cognitive maps (Behrens et al. 2018, Boorman et al. 2021). This region is strongly impacted by schizophrenia (Yasuda et al. 2022, Tamminga et al. 2010), and it communicates with PFC during goal-directed behavior (Schmidt, Duin, and Redish 2019). It is therefore possible that our results are driven by the interplay between representations in the hippocampal formation and goal-directed decision making mediated by DLPFC. Future neuroimaging studies can explicitly test this hippocampal-prefrontal relationship by simultaneously measuring map formation in the hippocampus, as well as its interactions with PFC during goal-directed behavior.

Our work has several limitations. First, because our data were collected as part of a larger initiative (Barch et al. 2023), we do not have access to patient medication status for the specific session in which they engaged in the tasks presented here, and so we are unable to control for this. Previous studies, however, have reported a lack of relationships between reinforcement learning and medication in schizophrenia (Culbreth et al., 2016; Geana et al., 2022). Concerning our specific variant of the two-step task, patients and controls may have learned the transition structure at different rates. However, in previous work, we have shown incremental learning of the transition structure these values quickly converge to the true structure regardless of the learning rate (Karagoz, Reagh, and Kool 2024). Finally, even though we used a guiding prompt during the behRSA task (“*In terms of the task, how related do you think these objects are?”*), it’s possible that patients and controls interpreted this task differently. Though we cannot rule out differences in basic task comprehension, we note that all participants were given an opportunity to ask for clarification pertaining to instructions as needed. Neuroimaging work, which would allow us to measure task representations implicitly, would ameliorate this concern.

In sum, our study shows that schizophrenia affects how people construct models of the environment. This may at least partly explain deficits in goal-directed control, which have long been considered a defining feature of the disorder. These results provide novel insights into the potential mechanisms underlying cognitive deficits in schizophrenia, and may inspire experimental manipulations that help patients construct better cognitive maps, which we predict will lead to increased goal-directed control. In turn, this could lead to specific, targeted interventions for the treatment of negative symptoms in schizophrenia.

## Declarations

## Availability of data and materials

Anonymized data for the decision-making task and the behavioral RSA task are available on the GitHub repository for the paper https://github.com/cdm-lab/sz-cog-maps-paper. Data for the working memory task, and self-reports is available on request.

## Code availability

Code used for the processing, analyses, and visualizations, are available on the GitHub repository for the paper https://github.com/cdm-lab/sz-cog-maps-paper

## Ethics approval

Approval was obtained from the Institutional Review Board of Washington University in St. Louis. The procedures used in this study adhere to the tenets of the Declaration of Helsinki.

## Consent to participate

Informed consent was obtained from all individual participants included in the study.

## Consent to publish

Not applicable.

## Author contributions

**Ata Karagoz:** Conceptualization, Methodology, Software, Formal Analyses, Writing – Original Draft, Writing – Review & Editing, Visualization. **Erin Moran:** Methodology, Investigation, Project administration, Data Curation, Writing – Review & Editing. **Deanna Barch:** Conceptualization, Resources, Funding Acquisition, Writing – Review & Editing. **Zachariah Reagh:** Conceptualization, Methodology, Writing – Review & Editing, Supervision. **Wouter Kool:** Conceptualization, Methodology, Writing – Review & Editing, Supervision.

## Acknowledgements

We would like to thank Jacob Pine and Nada Dalloul for helpful discussions. We would also like to thank Rachel Ryan for assistance in data collection. Finally, we would like to thank the members of the Complex Memory Lab, the Control and Decision Making Lab, and the Cognitive Control and Psychopathology lab at Washington University for their support.

