## Supplemental Materials for "Evidence for shallow cognitive maps in schizophrenia"

### Supplementary Methods

#### Reinforcement learning model

We adapted an established hybrid reinforcement learning model that we used in prior work to assess participants' behavior in the decision-making task, specifically dissociating model-free and model-based decision making. Every trial $t$ started out in one of two first-stage states $\left(s_{1,t}\right)$ where one of two possible actions $a_A$ and $a_B$ could be selected $\left(a_{1,t}\right)$. Depending on their selection, the participant deterministically transitioned to one of two second-stage states $\left(s_{2,t}\right)$ where they could perform only one action $\left(a_{2,t}\right)$ and then obtain a reward $(r_t)$. The model described here contains both a model-free learner and a model-based learner that learn expectations of long-term future reward $Q(s,a)$ for each combination of state and action. The model-free system learns reward expectations for each of the four teleporters and two generators, by updating their values based on reward prediction errors. The model-based system, on the other hand, learns a transition structure that represents to which planet each teleporter leads. It then combines this with the model-free reward expectations of the terminal, second-stage states to select between teleporters.

##### Model-free system

All model-free reward expectations were instantiated with a reward expectation of 4.5 (arithmetic mean of minimum and maximum possible reward) for all actions and states. The model-free learner would then use the $SARSA(\lambda)$ temporal difference learning algorithm to update its cached reward expectations based on the difference between predicted and received rewards. In the decision-making task this resulted in a reward prediction error $(\delta)$ being calculated at each stage according to:

$$\delta_{1,t} = Q_{MF}\left(S_{2,t}, a_{2,t}\right) - Q_{MF}\left(s_{1,t}, a_{1,t}\right)$$
$$\delta_{2,t} = r_t - Q_{MF}\left(s_{2,t}, a_{2,t}\right)$$

Notice that the second-stage prediction error incorporates the immediate reward outcome for that trial, but that the first-stage prediction error only incorporates expectations of future reward. The values of each prediction error were then used to update the reward expectations of the model-free learner at both the first and second stage:

$$Q_{MF}\left(s_{1,t}, a_{1,t}\right) \leftarrow Q_{MF}\left(s_{1,t}, a_{1,t}\right) + \alpha\delta_{1,t} + \alpha\lambda\delta_{2,t}$$
$$Q_{MF}\left(s_{2,t}, a_{2,t}\right) \leftarrow Q_{MF}\left(s_{2,t}, a_{2,t}\right) + \alpha\delta_{2,t}$$

Here, $\alpha$ is the reward learning rate (between 0 to 1) that determines how quickly new information about rewards is incorporated into the model-free learner expectations. The eligibility trace decay parameter $\lambda$ (between 0 to 1) determines how much a reward prediction error experienced after the second-stage choice changes first-stage reward expectations.

**Model-based system**

The model-based system uses the transition structure of the task to flexibly compute reward expectations for each available teleporter. Specifically, it has a transition matrix $T(s_1, a_1)$ that encodes the probability of moving to the second-stage state $s_2$ after choosing the action $a_1$ in the first-stage state $s_1$. In order to compute the model-based reward expectations, these probabilities are combined with the reward expectations at the second-stage:

$$Q_{MB}(s_{1,t}, a_{1,t}) = \sum_{s2} T\,(s_{1,t}, a_{1,t}) Q_{MB}(s_2, a_2)$$
$$Q_{MB}(s_{2,t}, a_{2,t}) = Q_{MF}(s_{2,t}, a_{2,t})$$

**Choice rule**

The model-free and model-based learners reward expectations in the first-stage states are integrated using a model-based weighting parameter $w$ (ranging from 0 to 1) using the following rule:

$$Q_{net}(s_1, a_1) = (1 - w) Q_{MF}(s_1, a_1) + w Q_{MB}(s_1, a_1)$$

We then used a softmax function to map the reward expectations to choice probabilities:

$$P(a_{1,t} = a_1 | s_{1,t}) = \frac{exp(\beta[Q_{net}(s_{1,t}, a_1)])}{\sum_a \; exp(\beta[Q_{net}(s_{1,t}, a)])}$$

Here, $\beta$ is the inverse softmax temperature (left-bounded to 0) that determines how much influence reward expectations have on choice probabilities and can be thought of as a measure of exploration and exploitation. High softmax temperatures mean that the model is more likely to explore and low softmax temperatures mean the model more commonly exploits its knowledge.

Due to the tendency of participants to perseverate on choices that are suboptimal we added two parameters to capture both response key and stimulus ‘stickiness’. The choice stickiness parameter $\pi$ (left unbounded) related to choice perseveration when positive and choice switching when negative. The response stickiness parameter $\rho$ captured perseveration of the response key press when positive and switching of response key press when negative.

$$rep(a_1) \quad = \begin{cases} 1 & \text{if } a_{1,t} = a_{1,t-1} \\ 0 & \text{otherwise.} \end{cases}$$
$$resp(a_1) \quad = \begin{cases} 1 & \text{if response for } a_{1,t} = \text{response for } a_{1,t-1} \\ 0 & \text{otherwise.} \end{cases}$$

With the addition of these perseveration parameters the full choice function is as follows:

$$P(a_{1,t} = a_1 | s_{1,t}) = \frac{exp(\beta[Q_{net}(s_{1,t}, a_1) + \pi * rep(a_1) + \rho * resp(a_1)])}{\sum_a \ exp(\beta[Q_{net}(s_{1,t}, a) + \pi * rep(a) + \rho * resp(a)])}$$

Together this results in a model with 6 free parameters which are fit using a *maximum a posteriori* (MAP) fitting procedure defined below.

**Model fitting procedure**

For each participant we obtained *maximum a posteriori* (MAP) estimates of the free parameters in the model, using custom scripts coupled with the 'scipy.optimize.minimize' function. All parameters had the following priors:

$$\begin{aligned} \alpha, \lambda, w &\sim \text{Beta}(2,2), \\ \beta &\sim \text{Gamma}(3,0.2), \\ \pi, \rho &\sim \mathcal{N}(0,1). \end{aligned}$$

These priors were empirically derived in work by Bolenz and colleagues (Bolenz et al. 2019). In order to avoid local optima, we randomly initialized the parameters and performed the optimization procedure 10 times per participant. We then selected the parameters of the run with the highest posterior probability.

To investigate the degree to which patients and controls altered their use of model-based control in response to motivational manipulations. We also fit a version of this model where a separate $w$ parameter was estimated for high-stakes and low-stakes trials. The difference between these parameters indicated the degree to which patients modulated their control in response to the stakes (Kool, Gershman, and Cushman 2017; Bolenz et al. 2019; Karagoz, Reagh, and Kool 2024).

### Supplementary Results

#### Points earned by stake in both groups:

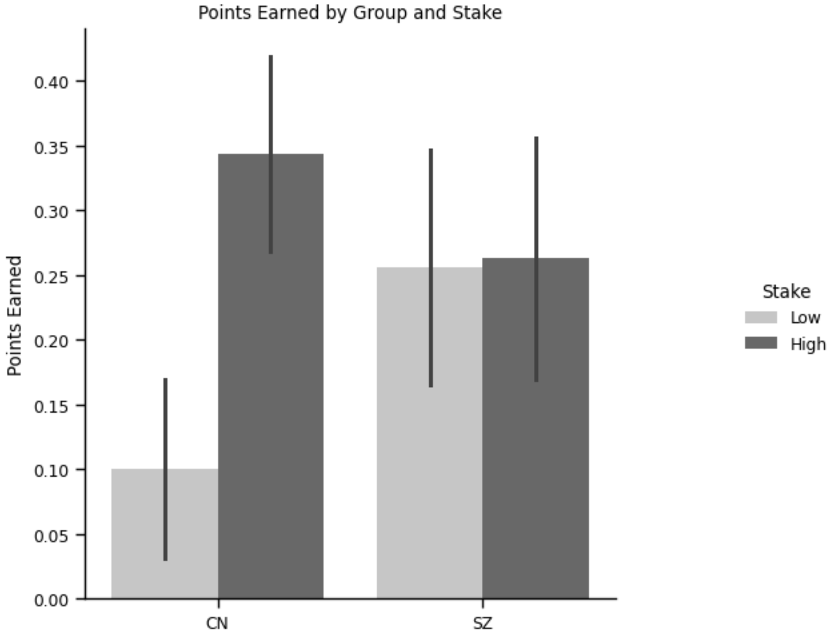

Figure S1: Points earned in the different stake condition for both the patients and controls. The data in this plot are the same to produce the plot Figure 2C. Errorbars are standard error of the mean.

#### Model-based control and performance with inverse temperature:

We found an interesting dissociation between performance and model-based control in our clinical sample. We hypothesized that this might be explained by the inverse temperature parameter ($\beta$) which governs the degree to which participants exploit their knowledge of rewards. Though we found no evidence of differences in the parameter between our control and clinical sample, we sought to investigate if there was a deeper moderative effect of $\beta$ on the relationship between $w$ and performance (as measured by points earned). To do this we ran a linear model of the following form:

Points Earned ~ $w$ x $\beta$ x Clinical Group

We found no evidence of moderation and when incorporating both $w$ and $\beta$ we no longer had a main effect of either.

| | Estimate | 2.5_ci | 97.5_ci | SE | DF | T-stat | P-val | Sig |
|---|---|---|---|---|---|---|---|---|
| Intercept | -0.286 | -0.962 | 0.389 | 0.333 | 35 | -0.86 | 0.396 | |
| $w$ | 0.551 | -0.68 | 1.782 | 0.606 | 35 | 0.909 | 0.37 | |
| $\beta$ | 0.168 | -0.211 | 0.546 | 0.187 | 35 | 0.898 | 0.375 | |

| | | | | | | | |
|---|---|---|---|---|---|---|---|
| $w : \beta$ | -0.076 | -0.715 | 0.563 | 0.315 | 35 | -0.241 | 0.811 |
| Clinical Group | -0.09 | -1.022 | 0.841 | 0.459 | 35 | -0.197 | 0.845 |
| $w$ : Clinical Group | 0.137 | -1.648 | 1.922 | 0.879 | 35 | 0.156 | 0.877 |
| $\beta$ : Clinical Group | 0.111 | -0.344 | 0.566 | 0.224 | 35 | 0.496 | 0.623 |
| $w$:$\beta$ : Clinical Group | -0.128 | -0.962 | 0.705 | 0.411 | 35 | -0.312 | 0.757 |

Table S1: Points Earned ~ $w$ x $\beta$ x Clinical Group estimates.

**Model-based control by behRSA**

In line with our prior work (Karagoz, Reagh, and Kool 2024), we hypothesized that planning relevant features such as the direct and indirect item associations would predict use of model-based control. We wondered if this effect differed across the clinical groups. To this end we ran a linear model of the following form:

$w$ ~(Visual Co-occurrence + Direct Item Association + Indirect Item Association x Clinical Group

We did not find a main effect either direct item or indirect item associations as we have previously reported (Karagoz, Reagh, and Kool 2024). There was also no evidence of an interaction.

| | Estimate | 2.5_ci | 97.5_ci | SE | DF | T-stat | P-val | Sig |
|---|---|---|---|---|---|---|---|---|
| Intercept | 0.549 | 0.438 | 0.659 | 0.055 | 35 | 10.063 | 0.0 | *** |
| Visual Co-occurrence | 0.003 | -0.002 | 0.008 | 0.003 | 35 | 1.132 | 0.265 | |
| Direct Item Association | 0.002 | -0.001 | 0.005 | 0.002 | 35 | 1.119 | 0.271 | |
| Indirect Item Association | -0.0 | -0.003 | 0.003 | 0.002 | 35 | -0.182 | 0.857 | |
| Clinical Group | -0.043 | -0.214 | 0.128 | 0.084 | 35 | -0.506 | 0.616 | |
| Visual Co-occurrence:Clinical Group | -0.002 | -0.008 | 0.005 | 0.003 | 35 | -0.474 | 0.638 | |
| Direct Item Association:Clinical Group | -0.005 | -0.01 | 0.001 | 0.003 | 35 | -1.608 | 0.117 | |

| | | | | | | | |
|---|---|---|---|---|---|---|---|
| Indirect Item Association:Clinical Group | 0.003 | -0.001 | 0.008 | 0.002 | 35 | 1.432 | 0.161 |

Table S2: *w* ~(Visual Co-occurrence + Direct Item Association +Indirect Item Association x Clinical Group

### MAP-SR and Snaith-Hamilton

Along with the associations reported in the main text we reasoned that hedonic capacity as measured by the Snaith-Hamilton, and motivation as measured by MAP-SR, would predict other features in the task. To this end we ran a series linear models looking at the effects of the above self-report measures on the modulation of control and performance in stakes conditions. We also wanted to assess whether either of these measures predicted participant cognitive maps.

**Snaith and MAP-SR on stakes control modulation:**

First we ran a linear model to assess the effect of self-report measures on the modulation of control in high vs low stakes contexts and to see whether this was different across clinical groups. We ran a model of the following form:

Difference in *w* ~ (Snaith-Hamilton + MAP-SR) x Clinical Group

We found no significant main effects for either of the two self-report measures as well as no significant interactions. This indicates that though increased hedonic capacity seems to be coupled with decreased use of model-based control in individuals with schizophrenia (as reported in the primary results), it does not predict their modulation of that control.

| | Estimate | 2.5_ci | 97.5_ci | SE | DF | T-stat | P-val | Sig |
|---|---|---|---|---|---|---|---|---|
| Intercept | 0.091 | 0.017 | 0.165 | 0.037 | 37 | 2.477 | 0.018 | * |
| Snaith-Hamilton | 0.023 | -0.066 | 0.113 | 0.044 | 37 | 0.526 | 0.602 | |
| MAP-SR | -0.023 | -0.099 | 0.053 | 0.038 | 37 | -0.619 | 0.54 | |
| Clinical Group | -0.058 | -0.163 | 0.047 | 0.052 | 37 | -1.111 | 0.274 | |
| Snaith-Hamilton:Clinical Group | -0.069 | -0.184 | 0.045 | 0.056 | 37 | -1.233 | 0.225 | |
| MAP-SR:Clinical Group | 0.077 | -0.03 | 0.185 | 0.053 | 37 | 1.465 | 0.151 | |

Table S3: Difference in *w* ~ (Snaith-Hamilton + MAP-SR) x Clinical Group

**Snaith and MAP-SR on stakes performance modulation:**

We also sought to assess whether the direct modulation of performance in the high compared to low stakes was linked to our self-report measures. To this end we ran a linear model of the following form:

Difference in *points* ~ (Snaith-Hamilton + MAP-SR) x Clinical Group

We found a significant main effect of the Snaith-Hamilton, indicating that higher hedonic response was coupled with more high-stakes enhancement of performance in controls. We also found a significant interaction with Snaith-Hamilton measure and clinical group such that increased hedonic response was not coupled with increased performance enhancement.

| | Estimate | 2.5_ci | 97.5_ci | SE | DF | T-stat | P-val | Sig |
|---|---|---|---|---|---|---|---|---|
| Intercept | 0.13 | -0.071 | 0.33 | 0.099 | 37 | 1.311 | 0.198 | |
| Snaith-Hamilton | 0.296 | 0.055 | 0.537 | 0.119 | 37 | 2.491 | 0.017 | * |
| MAP-SR | 0.072 | -0.134 | 0.277 | 0.101 | 37 | 0.708 | 0.483 | |
| Clinical Group | -0.099 | -0.382 | 0.184 | 0.14 | 37 | -0.707 | 0.484 | |
| Snaith-Hamilton:Clinical Group | -0.326 | -0.633 | -0.018 | 0.152 | 37 | -2.146 | 0.039 | * |
| MAP-SR:Clinical Group | 0.045 | -0.244 | 0.334 | 0.143 | 37 | 0.316 | 0.754 | |

Table S4: Difference in *points* ~ (Snaith-Hamilton + MAP-SR) x Clinical Group

**Snaith and MAP-SR on direct item association:**

After assessing the degree to which our self-report measures predicted modulations of task performance and control, we sought to assess their relationship with aspects of participant cognitive maps. First, we assessed whether our self-report measures were related to participant use of direct item representations. To do this we used a linear model of the following form:

Direct Item Association ~ (Snaith-Hamilton + MAP-SR) x Clinical Group

We found no effect of our self-report measures on the amount direct item association in participant cognitive maps.

| | Estimate | 2.5_ci | 97.5_ci | SE | DF | T-stat | P-val | Sig |
|---|---|---|---|---|---|---|---|---|
| Intercept | 16.43 | -0.95 | 33.809 | 8.577 | 37 | 1.915 | 0.063 | . |
| Snaith-Hamilton | 10.477 | -10.431 | 31.386 | 10.319 | 37 | 1.015 | 0.317 | |
| MAP-SR | -11.813 | -29.614 | 5.988 | 8.785 | 37 | -1.345 | 0.187 | |
| Clinical Group | -1.256 | -25.79 | 23.277 | 12.108 | 37 | -0.104 | 0.918 | |

| | | | | | | | |
|---|---|---|---|---|---|---|---|
| Snaith-Hamilton:Clinical Group | -10.054 | -36.719 | 16.612 | 13.16 | 37 | -0.764 | 0.45 |
| MAP-SR:Clinical Group | 3.216 | -21.832 | 28.264 | 12.362 | 37 | 0.26 | 0.796 |

Table S5: Direct Item Association ~ (Snaith-Hamilton + MAP-SR) x Clinical Group

**Snaith and MAP-SR on indirect item association:**

Next, we tested whether either self-report measure was related to indirect item associations using a linear model of the following form:

Indirect Item Association ~ (Snaith-Hamilton + MAP-SR) x Clinical Group

We found no main effects for either self-report measure, as well as a lack of interaction effects. This indicates that the self-report measures are potentially distinct from aspects of participant cognitive maps.

| | Estimate | 2.5_ci | 97.5_ci | SE | DF | T-stat | P-val | Sig |
|---|---|---|---|---|---|---|---|---|
| Intercept | 25.024 | 6.297 | 43.752 | 9.243 | 37 | 2.708 | 0.01 | * |
| Snaith-Hamilton | -0.155 | -22.685 | 22.376 | 11.12 | 37 | -0.014 | 0.989 | |
| MAP-SR | -14.794 | -33.976 | 4.387 | 9.467 | 37 | -1.563 | 0.127 | |
| Clinical Group | -25.375 | -51.812 | 1.062 | 13.047 | 37 | -1.945 | 0.059 | . |
| Snaith-Hamilton:Clinical Group | -2.953 | -31.686 | 25.781 | 14.181 | 37 | -0.208 | 0.836 | |
| MAP-SR:Clinical Group | 6.702 | -20.289 | 33.693 | 13.321 | 37 | 0.503 | 0.618 | |

Table S6: Indirect Item Association ~ (Snaith-Hamilton + MAP-SR) x Clinical Group

### Working Memory

For analyses reported in the following section, we have missing data from a single control participant, so those were not included. Thus, in the following section, we report analyses with 22 control participants and 20 individuals with schizophrenia.

**Model-based control and performance with working memory:**

First, we hypothesized that working memory capacity might be a moderating factor that was causing the dissociation between model-based control and performance in patients. To assess this possibility, we ran a model of the following form:

Points Earned ~ *w* x Running Span x Clinical Group

We found no main effects as well as no interaction effects indicating that working memory was not differentially moderating the relationship between *w* and performance in patients and controls.

| | Estimate | 2.5_ci | 97.5_ci | SE | DF | T-stat | P-val | Sig |
|---|---|---|---|---|---|---|---|---|
| Intercept | -0.377 | -1.479 | 0.724 | 0.542 | 34 | -0.696 | 0.491 | |
| *w* | 0.872 | -1.099 | 2.844 | 0.97 | 34 | 0.899 | 0.375 | |
| Running-span | 0.006 | -0.021 | 0.034 | 0.013 | 34 | 0.488 | 0.629 | |
| *w :* Running-span | -0.008 | -0.05 | 0.035 | 0.021 | 34 | -0.355 | 0.725 | |
| Clinical Group | 1.123 | -0.226 | 2.472 | 0.664 | 34 | 1.691 | 0.1 | . |
| *w* : Clinical Group | -1.761 | -4.216 | 0.694 | 1.208 | 34 | -1.458 | 0.154 | |
| Running-span : Clinical Group | -0.021 | -0.053 | 0.012 | 0.016 | 34 | -1.306 | 0.2 | |
| *w* : Running-span : Clinical Group | 0.035 | -0.022 | 0.093 | 0.028 | 34 | 1.248 | 0.22 | |

Table S7: Points Earned ~ *w* x Running Span x Clinical Group

**Direct item association predicted by working memory capacity:**

We next sought to assess the effects of working memory capacity on both of the planning-relevant features of participants' cognitive maps. Both the direct item association and indirect item association require integration of events over time and so we reasoned that these should be predicted by working memory capacity. First, we assessed whether working memory capacity (as measured by the running span), predicted the presence of direct item association in participant cognitive maps. We used a linear model of the following form:

Direct Item Association ~ (Running Span) x Clinical Group

We found a significant main effect for running span indicating that increased working memory capacity was coupled with increased representation of direct item association in the cognitive maps of controls. We also found a significant interaction where this relationship was not present in individuals with schizophrenia. The results of this model can be seen in Figure S1.

| | Estimate | 2.5_ci | 97.5_ci | SE | DF | T-stat | P-val | Sig |
|---|---|---|---|---|---|---|---|---|
| Intercept | -49.074 | -97.853 | -0.295 | 24.095 | 38 | -2.037 | 0.049 | * |
| Running Span | 1.41 | 0.438 | 2.382 | 0.48 | 38 | 2.937 | 0.006 | ** |
| Clinical Group | 66.455 | 5.18 | 127.73 | 30.268 | 38 | 2.196 | 0.034 | * |
| Running Span:Clinical Group | -1.404 | -2.797 | -0.011 | 0.688 | 38 | -2.04 | 0.048 | * |

Table S8: Direct Item Association ~ (Running Span) x Clinical Group

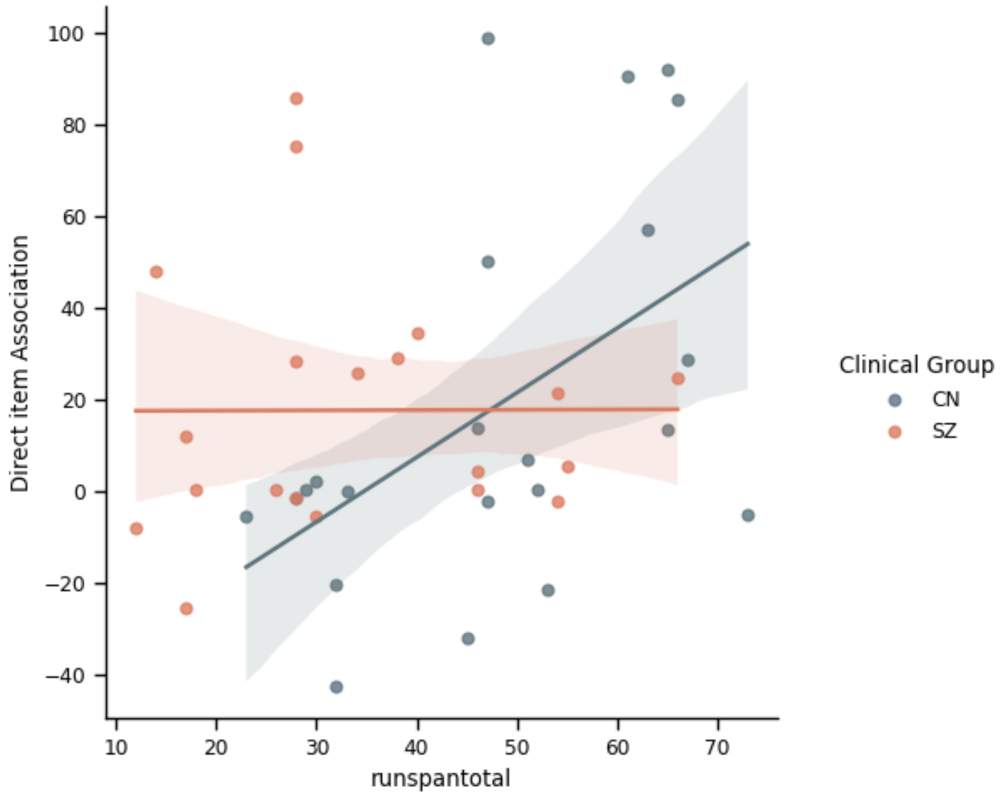

Figure S2: Correlations of Direct item association measure and working memory capacity by clinical group. There is a positive relationship between working memory and the direct item association measure in controls but not in individuals with schizophrenia.

**Indirect item association predicted by working memory capacity:**

We reasoned that due to the nature of requiring abstraction over a full set of trials, indirect item association would be predicted by working memory capacity. To test this, we ran a model of the form:

Indirect Item Association ~ (Running Span) x Clinical Group

Surprisingly, we found no effect of working memory capacity on the representation of indirect item associations in either our control or clinical groups.

| | Estimate | 2.5_ci | 97.5_ci | SE | DF | T-stat | P-val | Sig |
|---|---|---|---|---|---|---|---|---|
| Intercept | -5.251 | -62.323 | 51.821 | 28.192 | 38 | -0.186 | 0.853 | |
| Running Span | 0.593 | -0.545 | 1.73 | 0.562 | 38 | 1.055 | 0.298 | |
| Clinical Group | 7.421 | -64.272 | 79.114 | 35.415 | 38 | 0.21 | 0.835 | |
| Running Span:Clinical Group | -0.562 | -2.192 | 1.069 | 0.805 | 38 | -0.697 | 0.49 | |

Table S8: Indirect Item Association ~ (Running Span) x Clinical Group

### Differences by sex

We further wondered whether any of the results reported in the main text differed as a result of participant sex. We reran a series of models incorporating sex as a binary regressor with males coded as 0 and females coded as 1.

**Sex differences in behRSA:**

First, we reran our hierarchical mixed effects model with sex as an additional regressor.

Formula: Coefficient ~ complexity x Clinical Group x Sex + (1|subid)

We found no main effects of complexity, clinical group, or sex. We did find a trending effect between complexity and clinical group. This is perhaps difficult to interpret due to the small group sizes entering the model (approximately 10 participants for each sex x clinical group bin).

| | Estimate | 2.5_ci | 97.5_ci | SE | DF | T-stat | P-val | Sig |
|---|---|---|---|---|---|---|---|---|
| (Intercept) | 22.128 | 7.784 | 36.472 | 7.318 | 38.0 | 3.024 | 0.004 | ** |
| complexity | 8.302 | -2.36 | 18.964 | 5.44 | 80.0 | 1.526 | 0.131 | |
| Clinical Group | -8.228 | -28.968 | 12.513 | 10.582 | 38.0 | -0.777 | 0.442 | |
| Sex | -14.105 | -34.846 | 6.636 | 10.582 | 38.0 | -1.333 | 0.19 | |
| complexity: Clinical Group | -14.018 | -29.435 | 1.399 | 7.866 | 80.0 | -1.782 | 0.079 | . |
| complexity:Sex | -3.973 | -19.39 | 11.445 | 7.866 | 80.0 | -0.505 | 0.615 | |

| | | | | | | | |
|---|---|---|---|---|---|---|---|
| Clinical Group:Sex | 11.991 | -19.046 | 43.027 | 15.835 | 38.0 | 0.757 | 0.454 |
| complexity:Clinical Group:Sex | 2.532 | -20.538 | 25.602 | 11.771 | 80.0 | 0.215 | 0.83 |

Table S9: Coefficient ~ complexity x Clinical Group x Sex+(1|subid)

**Points earned ~ *w* x Clinical Group x Sex:**

We next focused on whether the dissociation between patient model-based control and performance could be partially explained by participant sex. To this end we ran a model of the form:

Points earned ~ *w* x Clinical Group x Sex

We found no main effects or interaction effects.

| | Estimate | 2.5_ci | 97.5_ci | SE | DF | T-stat | P-val | Sig |
|---|---|---|---|---|---|---|---|---|
| Intercept | -0.079 | -0.593 | 0.435 | 0.253 | 34 | -0.311 | 0.758 | |
| *w* | 0.523 | -0.237 | 1.284 | 0.374 | 34 | 1.398 | 0.171 | |
| Clinical Group | 0.076 | -0.611 | 0.764 | 0.338 | 34 | 0.226 | 0.823 | |
| *w*:Clinical Group | 0.218 | -0.931 | 1.368 | 0.566 | 34 | 0.386 | 0.702 | |
| Sex | -0.191 | -1.013 | 0.632 | 0.405 | 34 | -0.471 | 0.641 | |
| *w*:Sex | 0.329 | -1.023 | 1.68 | 0.665 | 34 | 0.494 | 0.625 | |
| Clinical Group:Sex | 0.491 | -0.615 | 1.597 | 0.544 | 34 | 0.902 | 0.374 | |
| *w*:Clinical Group:Sex | -1.419 | -3.303 | 0.466 | 0.927 | 34 | -1.529 | 0.135 | |

Table S10: Points earned ~ *w* x Clinical Group x Sex

***w* ~ Working Memory x Clinical Group x Sex**

We next wondered whether the dissociation in patients use of model-based control and their working memory capacity was accounted for by sex differences. We ran the following model:

*w* ~ Running-span x Clinical Group x Sex

We found a significant main effect of running span, indicating that in male controls it was coupled with increased model-based control. We found a significant main effect for clinical group, and a significant interaction where working memory capacity was not coupled with increased use of model-based control in patients. We also found a trending interaction between sex and working memory capacity indicating that working memory was less of a predictor of *w* in female controls than male controls. We also found a trending interaction between sex, working memory capacity, and clinical group. The results of this linear model can be more easily seen in Figure S2. The correlation of model-based control and working memory capacity seems to be primarily driven by male control participants.

| | Estimate | 2.5_ci | 97.5_ci | SE | DF | T-stat | P-val | Sig |
|---|---|---|---|---|---|---|---|---|
| Intercept | 0.071 | -0.322 | 0.464 | 0.193 | 33 | 0.369 | 0.715 | |
| Running Span | 0.012 | 0.004 | 0.019 | 0.004 | 33 | 3.067 | 0.004 | ** |
| Clinical Group | 0.579 | 0.08 | 1.078 | 0.245 | 33 | 2.36 | 0.024 | * |
| Running Span:Clinical Group | -0.017 | -0.029 | -0.005 | 0.006 | 33 | -2.82 | 0.008 | ** |
| Sex | 0.481 | -0.101 | 1.063 | 0.286 | 33 | 1.683 | 0.102 | |
| Running Span:Sex | -0.011 | -0.023 | 0.001 | 0.006 | 33 | -1.943 | 0.061 | . |
| Clinical Group:Sex | -0.608 | -1.375 | 0.159 | 0.377 | 33 | -1.612 | 0.117 | |
| Running Span:Clinical Group:Sex | 0.017 | -0.001 | 0.036 | 0.009 | 33 | 1.937 | 0.061 | . |

Table S11: *w* ~ Running-span x Clinical Group x Sex

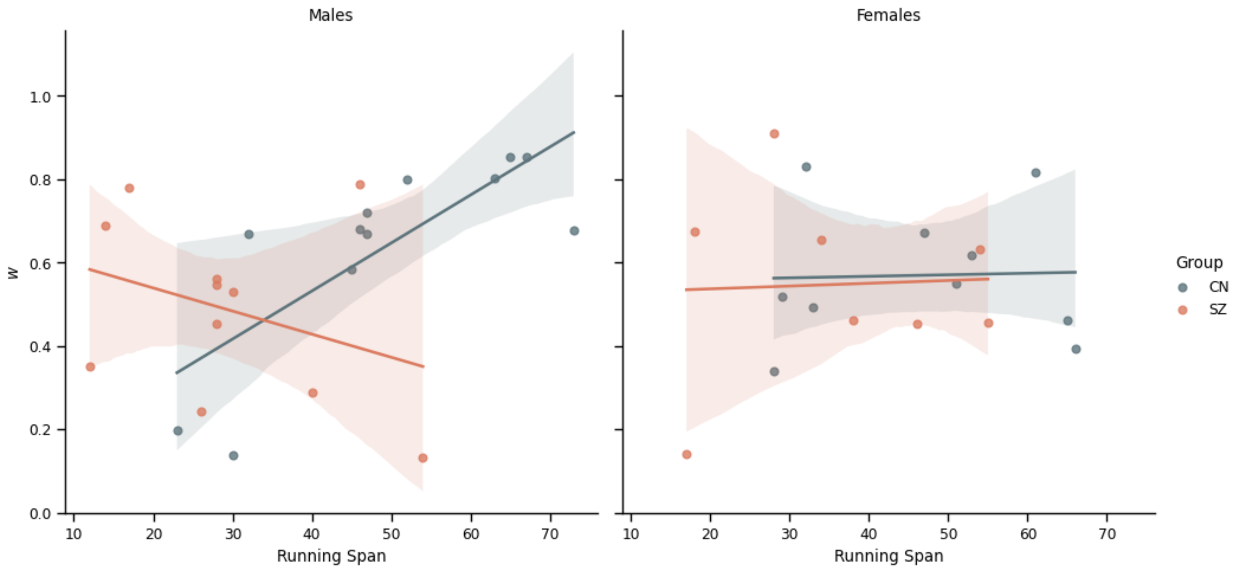

Figure S2: Differences in associations between working memory (as measured by running span) and model-based control in patient and control groups divided by sex.

***w* ~ Snaith-Hamilton + MAP-SR x Clinical Group x Sex:**

We further wanted to assess whether the use of model-based control we dependent on our self-report data, and whether this differed by participant sex.

We ran a model of the form: *w* ~ Snaith-Hamilton + MAP-SR x Clinical Group x Sex

We found a main effect in the negative direction for the MAP-SR indicating that increased motivation was correlated with decreased use of model-based control in male control participants. We also found a main effect of group, as well as a significant interaction indicating that males with schizophrenia had a positive relationship between use of model-based control and their self-reported motivation. We found a significant interaction between MAP-SR and Sex such that control females had a positive relationship between motivation and use of control. Finally, we found a three-way interaction of MAP-SR, clinical group, and sex. This indicates that females with schizophrenia have a negative relationship between their motivation as measured by the MAP-SR and use of model-based control. These findings are difficult to interpret given the small sizes for each group.

| | Estimate | 2.5_ci | 97.5_ci | SE | DF | T-stat | P-val | Sig |
|---|---|---|---|---|---|---|---|---|
| Intercept | 0.671 | 0.556 | 0.786 | 0.056 | 30 | 11.938 | 0.0 | *** |
| Snaith-Hamilton | 0.062 | -0.061 | 0.184 | 0.06 | 30 | 1.029 | 0.312 | |
| MAP-SR | -0.159 | -0.278 | -0.04 | 0.058 | 30 | -2.739 | 0.01 | * |
| Clinical Group | -0.207 | -0.372 | -0.042 | 0.081 | 30 | -2.565 | 0.016 | * |
| Snaith-Hamilton:Clinical Group | -0.13 | -0.305 | 0.044 | 0.085 | 30 | -1.528 | 0.137 | |
| MAP-SR:Clinical Group | 0.285 | 0.109 | 0.462 | 0.086 | 30 | 3.301 | 0.002 | ** |
| Sex | -0.121 | -0.289 | 0.048 | 0.083 | 30 | -1.46 | 0.155 | |
| Snaith-Hamilton:Sex | -0.05 | -0.228 | 0.128 | 0.087 | 30 | -0.571 | 0.572 | |
| MAP-SR:Sex | 0.186 | 0.017 | 0.355 | 0.083 | 30 | 2.249 | 0.032 | * |
| Clinical Group:Sex | 0.177 | -0.073 | 0.428 | 0.123 | 30 | 1.445 | 0.159 | |
| Snaith-Hamilton:Clinical Group:Sex | 0.039 | -0.277 | 0.355 | 0.155 | 30 | 0.252 | 0.802 | |
| MAP-SR:Clinical Group:Sex | -0.347 | -0.661 | -0.033 | 0.154 | 30 | -2.254 | 0.032 | * |

**Modulation of performance ~ behRSA x Clinical Group x Sex:**

Finally, we sought to assess whether the enhancement of performance in high stakes compared to low stakes trials was predicted differentially by the behRSA parameters in different clinical groups as well as by participant sex.

We fit a model of the form: Difference in points ~ (Visual Co-Occurrence + Direct Item Association + Indirect Item Association) x Clinical Group x Sex

We found a trending main effect of direct item association, indicating that the presence of that feature was coupled with increased enhancement of performance in high-stakes contexts for male controls. We also found a significant interaction effect for direct item association and clinical group such that males with schizophrenia did not show the relationship between increased representation of direct item associations and increased performance enhancement. Finally, we found a trending main effect for sex indicating that females had higher performance enhancement for high-stakes contexts than males.

| | Estimate | 2.5_ci | 97.5_ci | SE | DF | T-stat | P-val | Sig |
|---|---|---|---|---|---|---|---|---|
| Intercept | -0.136 | -0.542 | 0.271 | 0.198 | 26 | -0.687 | 0.498 | |
| Visual Co-occurrence | 0.009 | -0.006 | 0.025 | 0.008 | 26 | 1.214 | 0.236 | |
| Direct Item Association | 0.008 | -0.001 | 0.017 | 0.004 | 26 | 1.887 | 0.07 | . |
| Indirect Item Association | -0.002 | -0.01 | 0.006 | 0.004 | 26 | -0.562 | 0.579 | |
| Clinical Group | 0.037 | -0.482 | 0.555 | 0.252 | 26 | 0.145 | 0.886 | |
| Visual Co-occurrence:Clinical Group | 0.001 | -0.022 | 0.024 | 0.011 | 26 | 0.081 | 0.936 | |
| Direct Item Association:Clinical Group | -0.017 | -0.033 | 0.0 | 0.008 | 26 | -2.052 | 0.05 | . |
| Indirect Item Association:Clinical Group | 0.009 | -0.007 | 0.025 | 0.008 | 26 | 1.182 | 0.248 | |
| Sex | 0.477 | -0.009 | 0.963 | 0.236 | 26 | 2.018 | 0.054 | . |
| Visual Co-occurrence:Sex | 0.0 | -0.024 | 0.025 | 0.012 | 26 | 0.041 | 0.968 | |
| Direct Item Association:Sex | 0.004 | -0.012 | 0.019 | 0.008 | 26 | 0.483 | 0.633 | |
| Indirect Item Association:Sex | -0.007 | -0.02 | 0.007 | 0.007 | 26 | -0.992 | 0.33 | |
| Clinical Group:Sex | -0.151 | -1.023 | 0.721 | 0.424 | 26 | -0.356 | 0.725 | |

| | | | | | | | |
|---|---|---|---|---|---|---|---|
| Visual Co-occurrence:Clinical Group:Sex | -0.009 | -0.041 | 0.024 | 0.016 | 26 | -0.551 | 0.586 |
| Direct Item Association:Clinical Group:Sex | -0.007 | -0.056 | 0.042 | 0.024 | 26 | -0.29 | 0.774 |
| Indirect Item Association:Clinical Group:Sex | 0.005 | -0.019 | 0.03 | 0.012 | 26 | 0.438 | 0.665 |
